# De-glycosylated non-structural protein 1 enhances dengue virus clearance by limiting PD-L1/PD-1 mediated T cell apoptosis

**DOI:** 10.1101/2024.01.08.574590

**Authors:** Fakhriedzwan Idris, Donald Heng Rong Ting, Eunice Tze Xin Tan, Corrine Wan, Kuan Rong Chan, Peter I Benke, Jan Kazimierz Marzinek, Jack M Copping, Qin Hui Li, Ian Walsh, Jane R Allison, Peter John Bond, Federico Torta, Terry Nguyen-Khuong, Sylvie Alonso

**Affiliations:** Infectious Diseases Translational Research Programme, Department of Microbiology and Immunology, Yong Loo Lin School of Medicine, National University of Singapore, Singapore; Immunology Programme, Life Sciences Institute; National University of Singapore; Analytical Science & Technology (GlycoAnalytics), Bioprocessing Technology Institute, Agency for Science, Technology and Research A*STAR, Singapore; Emerging Infectious Diseases programme, Duke-NUS Medical School, Singapore; Precision Medicine Translational Research Programme and Department of Biochemistry, Yong Loo Lin School of Medicine, National University of Singapore, Singapore; Singapore Lipidomics Incubator, Life Sciences Institute, National University of Singapore, Singapore; Bioinformatics Institute, Agency for Science, Technology and Research A*STAR, Singapore; Biomolecular Interaction Centre, Digital Life Institute, Maurice Wilkins Centre for Molecular Biodiscovery, University of Auckland, Auckland, New Zealand; School of Biological Sciences, University of Auckland, Auckland, New Zealand; Department of Biological Sciences, National University of Singapore, Singapore

## Abstract

The non-structural protein 1 (NS1) of dengue virus (DENV) contains two highly conserved N-glycosylation sites at positions 130 and 207 (N130 and N207). Intracellular NS1 monomers and homo-dimers participate in viral RNA replication within membrane-bound replication complexes. Soluble multimeric NS1 (sNS1) is secreted into the extracellular milieu and represents an important virulence factor for DENV through its ability to interfere with the host complement activation cascade and to induce vascular leakage. The role of the two N-glycans in NS1 biological activities, however, has not been carefully examined. Here, stable DENV2 mutants that lack glycan at either N sites of NS1 were engineered. We showed that the lack of glycans at either N site of NS1 did not impair viral replication nor viral output in both mosquito and mammalian cell lines. In contrast, while N130 de-glycosylated DENV displayed parental *in vivo* fitness in IFNAR^-/-^ mice, the N207 de-glycosylated mutant was significantly attenuated as evidenced by 100% survival rate, which correlated with accelerated viral clearance in circulation. sNS1-depletion, sNS1 exogenous administration and co-infection experiments supported that N207 de-glycosylated NS1 exerted a dominant attenuating effect during *in vivo* infection. Bulk RNAseq, inflammatory cytokine profile, immune phenotyping of neutrophils and T cells, immune cell depletion and immune checkpoint blockade approaches led us to propose that N207 de-glycosylated NS1 limited CD8^+^ T cell apoptosis mediated by the PD-L1/PD-1 axis, thereby improving viral clearance efficacy. This work uncovers a novel immune evasion strategy where N207 glycans on NS1 prevent the protein from exerting immune modulation activity that would be detrimental to DENV.

## INTRODUCTION

Dengue poses a significant threat in the tropical and subtropical regions, impacting approximately 390 million individuals across more than 125 countries (Bhatt et al., 2013). Projections indicate that by 2080, this number could rise to 2.25 billion people (Messina et al., 2019). In these endemic countries, dengue is a major public health concern due to the limited efficacy of existing dengue vaccines, and the lack of specific antiviral treatment (Idris, Ting, & Alonso, 2021). Symptomatic dengue causes flu-like symptoms with fever and rash. However, it can develop into severe illness known as dengue haemorrhagic fever (DHF), characterised by coagulopathy, vascular leakage, and thrombocytopenia, which may further progress into dengue shock syndrome (DSS) (Martina, Koraka, & Osterhaus, 2009). The infection is caused by the mosquito-borne flavivirus, dengue virus (DENV) that has been classified under four antigenically distinct serotypes, DENV1-4 that co-circulate with largely unpredictable dominance patterns. Like other flaviviruses, DENV genome is a positive-sense, single stranded RNA of approximately 10.7 kb in size and encodes three structural and seven non-structural (NS) proteins. Among which, NS1 is highly conserved across DENV1-4 and comprises of 352 amino acids with a molecular weight ranging from 40 to 50 kDa, depending on its glycosylation status (Flamand et al., 1999). NS1 monomers and dimers are found intracellularly, with the dimers often membrane-associated, while the secreted, soluble form of NS1 (sNS1) has been described as an atypical barrel-shaped hexamer with a central lipid cargo (Gutsche et al., 2011). A recent study however has challenged this model and reported that sNS1 exists predominantly in either ‘loose’ or ‘stable’ tetrameric form with differential susceptibility to antibodies (Shu et al., 2022).

While the main role of intracellular NS1 is to support DENV genome replication (Zhang et al., 2023), *in vitro* and *in vivo* studies have reported a variety of biological activities for secreted sNS1, including interfering with complement activation pathways (M. F. Lee, Voon, Lim, Chua, & Poh, 2022) and inducing endothelial hyperpermeability. The latter was found to involve shedding of the glycocalyx lining the intravascular endothelial layer (Biering et al., 2021; Puerta-Guardo et al., 2019) and/or Toll-like receptor 4 (TLR4)-mediated production of pro-inflammatory cytokines (Modhiran et al., 2015). Other studies have also shown that sNS1 is highly immunogenic (Freire et al., 2017; Stettler et al., 2016), and that anti-NS1 antibodies afforded protection against severe disease in animal models (Bailey et al., 2019; Beatty et al., 2015; Gonçalves et al., 2015; Richner et al., 2017). However, recent *in vivo* studies showed that sNS1 was dispensable for highly virulent DENV2 strains whereby NS1-specific immunity was not protective (P. X. Lee et al., 2020; Lin et al., 2021).

DENV NS1 has two N-linked glycans at N130 and N207 position, which are conserved across the four DENV serotypes, and whose sugar composition is site-specific and depends on the cell type in which NS1 protein is produced (Thiemmeca et al., 2016). DENV NS1 produced in mammalian cells typically displays complex glycans at N130, whereas N207 sugars mainly consist of high-mannose (Pryor & Wright, 1994). These N-glycans have been proposed to confer stability to the NS1 dimer and to be involved in the secretion of multimeric sNS1 (Somnuke, Hauhart, Atkinson, Diamond, & Avirutnan, 2011; Thiemmeca et al., 2016). Moreover, abrogation of N130 or both N-glycosylation sites on NS1 resulted in reduced virus growth *in vitro* (Crabtree, Kinney, & Miller, 2005; Pryor & Wright, 1994; Tajima, Takasaki, & Kurane, 2008). On the other hand, N207 glycans were found to play an essential role in sNS1-induced glycocalyx layer disruption in endothelial cell cultures (Wang et al., 2019). *In vivo* studies reported that lack of glycans on NS1 at either site led to significant attenuation of DENV neurovirulence in suckling mice (Crabtree et al., 2005; Fang et al., 2023). The authors speculated that lower sNS1 levels in circulation were responsible for the observed attenuated phenotype, although no experimental evidence was provided to support this hypothesis.

Here, we investigated the role of glycans on NS1 produced by a non-mouse adapted DENV2 strain, namely D2Y98P, that belongs to the Cosmopolitan genotype. D2Y98P is a representative of DENV2 strains that circulate in Southeast Asia and that have often been involved in dengue outbreaks (Lee et al, 2021). In sharp contrast with studies using less fit DENV strains, our previous work has shown that in D2Y98P-infected Type I-interferon-deficient mice (IFNAR^-/-^), sNS1 does not contribute to the virus fitness and virulence, despite high levels of sNS1 in circulation (Lee et al, 2021). This unique feature prompted us to further investigate the role of glycans on NS1 in the context of D2Y98P infection. We postulated that those highly conserved N-glycans may still play an important role in DENV intrinsic fitness, as part of NS1’s role in the virus intracellular replication. However, our findings did not support this hypothesis, and instead unraveled a novel immune evasion strategy mediated by NS1 N207 glycans.

## RESULTS

### Production and characterization of de-glycosylated NS1 DENV mutants

DENV mutants that lack the glycan motifs on either N130 or N207 sites were generated by substituting one of the amino acids that make up the glycosylation motif N-X-S/T, where X represents any amino acid except Proline. All six mutants were viable and displayed plaque morphology that was comparable to parental D2Y98P strain (WT), except T209L NS1 mutant which displayed significantly smaller plaques (Suppl. Fig. S1). To evaluate the mutation stability, the virus mutant strains were serially passaged in mosquito C6/36 cells and their genome was sequenced. Mutants harboring N130Q and T209L substitutions were found stable after 3 passages in C6/36 cells, while the rest reverted to parental glycosylation motif (Suppl. Table S1). This observation thus indicated that the nature of the amino acid substitution within the glycosylation motif plays an important role in virus viability.

### N130Q NS1 and T209L NS1 DENV mutants display parental *in vitro* fitness, while T209L NS1 DENV is attenuated in mice

The *in vitro* fitness of N130Q and T209L NS1 DENV mutants in mosquito and mammalian cell lines was comparable to the WT as evidenced by similar viral titers (Fig. 1A) and soluble NS1 (sNS1) levels (Fig. 1B) measured in the culture supernatant. These results thus indicated that either NS1 glycans are dispensable for DENV replication and NS1 secretion *in vitro*. We next examined the role of NS1 glycosylation in DENV fitness in a symptomatic mouse model previously established in our laboratory (P. X. Lee et al., 2020). IFNAR^-/-^ mice infected with N130Q NS1 mutant displayed clinical manifestations, body weight loss profile and viremia titers that were comparable to mice infected with WT virus (Fig. 1C-F). In contrast, mice infected with T209L NS1 mutant displayed reduced disease severity as evidenced by milder clinical scores, limited body weight loss and 100% survival (Fig. 1 C-E). Furthermore, while the initial viremia profile and peak viremia titer (day 3 p.i.) in mice infected with T209L NS1 mutant were similar to those measured in WT-infected animals, lower titers were measured at days 5 and 6 p.i. (Fig. 1F), likely explaining the attenuated phenotype observed. Interestingly, lower systemic levels of sNS1 were measured in mice infected with either mutant compared to WT control (Fig. 1G), therefore ruling out the hypothesis that lower concentrations of circulating sNS1 accounts for the attenuated disease severity upon infection with T209L mutant. Together, these results indicated that glycans on NS1 at N207, but not at N130, play a critical role in D2Y98P strain fitness and virulence in IFNAR^-/-^ mice.

**Figure 1.**
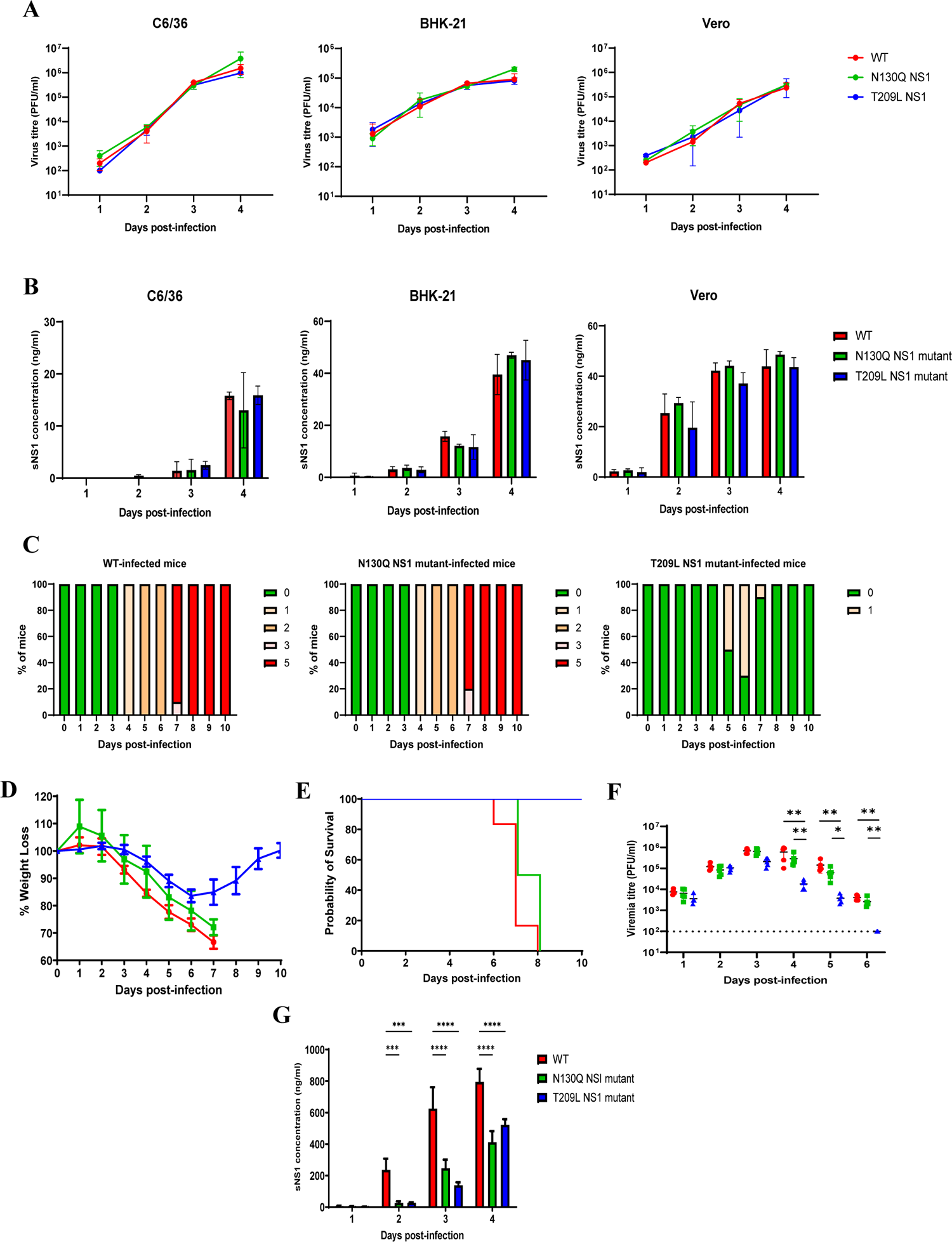
*In vitro* and *in vivo* fitness of N130Q and T209L NS1 mutants. A) Viral titers in culture supernatants. C6/36, BHK-21, and Vero cells were infected at MOI 0.1 with D2Y98P (WT), N130Q or T209L NS1 mutants. Viral titers in the culture supernatants were measured by plaque assay and were expressed as mean ± SD of triplicates. The detection limit was set at 10^2^ pfu/ml. B) sNS1 levels in culture supernatant. sNS1 concentrations were determined by ELISA. Results were expressed as mean ± SD of triplicates. C-G) IFNAR^-/-^ mice were infected with 10exp6 PFU of each virus via the subcutaneous (sc.) route. C-E) Survival, body weight loss profile and clinical scores (n=10); 0 – healthy, 1 – ruffled fur, 2 – hunched back, 3 – lethargy, 4 – limb paralysis, 5 – mice displaying 30% weight loss (euthanasia). F) Viremia titers were determined at the indicated points by plaque assay (n=5). G) Systemic sNS1 levels were measured by ELISA (n=5). Data shown are representative of at least 2 independent biological repeats.

### Monomeric and multimeric T209L NS1 display 3D structure, conformation and lipid cargo composition comparable to WT NS1

Since T209L substitution on NS1 was found to significantly impact DENV *in vivo* fitness, we hypothesized that lack of the glycans at N207 may disrupt the conformation and/or lipid composition of NS1 hexamer, that may in turn influence its biological properties. First, comparable 3D structure of WT and T209L NS1 monomers was obtained from AlphaFold2 (Fig. 2A). Second, when purified WT and T209L NS1 proteins were analyzed in native PAGE followed by Western blot, a comparable high molecular weight single band (corresponding to multimeric NS1) was detected, although T209L NS1 migrated slightly further than WT NS1, likely due to the lack of glycans at position 207 (Fig. 2B). Third, comparable lipid cargo composition was detected by mass spectrometry, which consisted of six dominant lipid classes, namely phosphatidylcholine, phosphatidylethanolamine, phosphatidylinositol, sphingomyelin, GM3 and hexosylceramide species, as previously described (Gutsche et al. 2011) (Fig. 2C). A reduction in some of sphingomyelin and GM3 species was however noticeable with T209L NS1 protein (Fig. 2C). Finally, all atom molecular dynamics (MD) simulations were performed in explicit water of NS1 WT or T209L NS1 and their lipid cargo. The glycan motifs at N130 and N207 used in these simulations were those experimentally determined by glycoproteomics (Suppl. Fig. S2). Simulations with WT NS1 indicated many interactions between N130 glycans localized away from the hydrophobic core of the hexamer, near the edges of the protein where lipids could potentially dissociate (Fig. 2D). In contrast glycans on N207 were positioned lower with respect to the hexamer center of mass near the hydrophobic core domain (residues 1-29), with negligible interactions with lipids (Fig. 2D). Therefore, N130 glycans appeared to keep the lipids inside of the hexameric structure. Simulation with T209L NS1 led to similar findings whereby N130 sugars displayed many contacts with lipids which were well-contained inside the hexamer (Fig. 2D).

**Figure 2.**
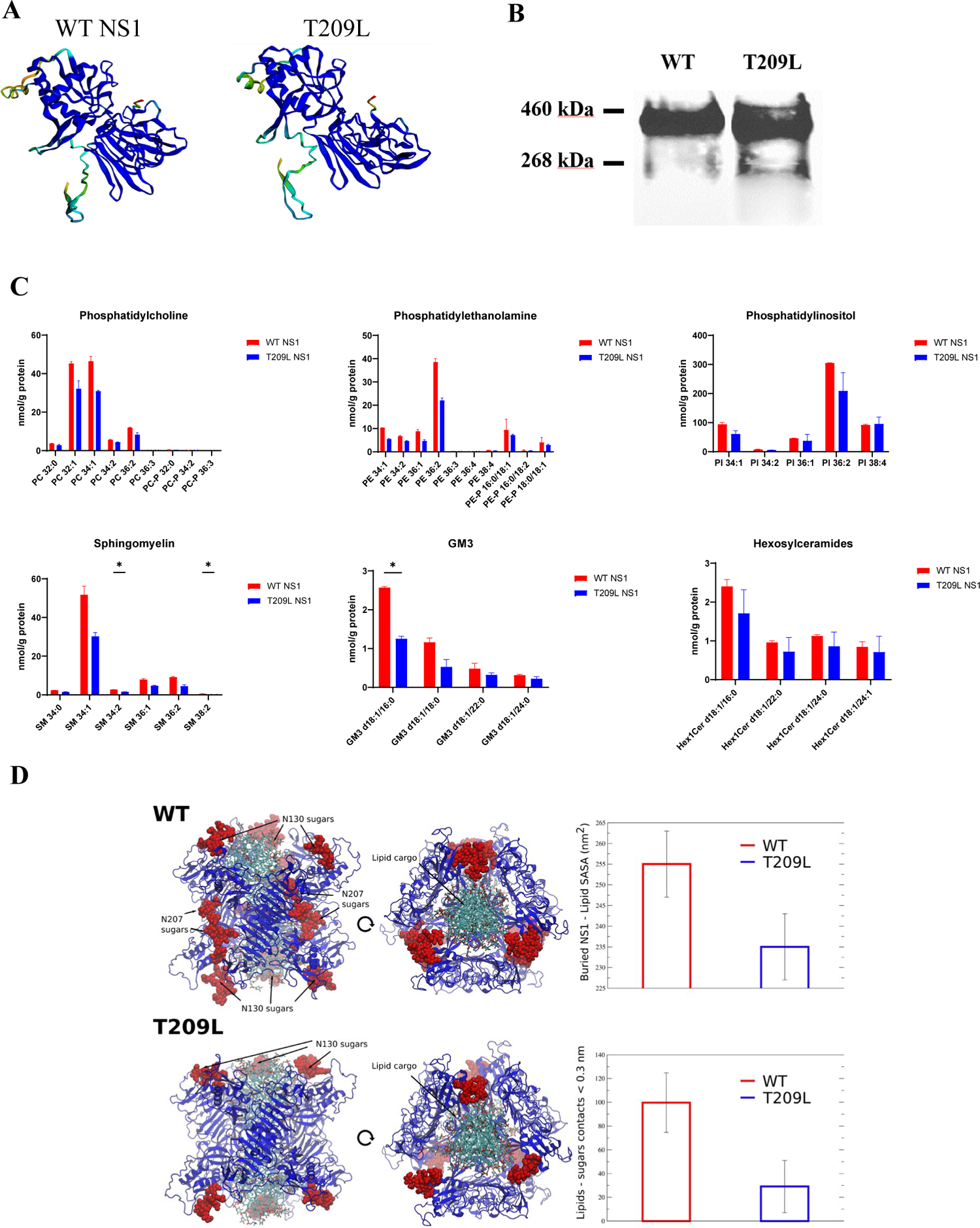
Structural and biochemical features of purified WT and T209L NS1 proteins. A) Simulated 3D structure of monomeric WT and T209L NS1 proteins using AlphaFold2 software. B) Detection of multimeric T209L and WT sNS1 by native PAGE and Western blot. C) Mass spectrometry analysis of the lipid composition of WT and T209L sNS1 proteins. D) Representative snapshots from MD simulations of WT and NS1 mutants. Cartoon representation from side (left) and top (right) views. NS1 protein is shown in blue. Lipids are shown in licorice representation (cyan-carbon, blue-nitrogen, red-oxygen) while glycans are shown as red spheres. The graphs shows last 100 ns averaged number of contacts between lipids and glycans as well as NS1:glycan – lipid buried SASA. The buried solvent accessible surface area (SASA) between NS1:glycans and lipid cargo was calculated as a sum of NS1:glycans and lipids before subtracting (TLR4:glycan)–lipid SASA. The error bars represent standard deviation (SD). Data shown are representative of at least 2 independent repeats.

Together, these observations suggested that the T209L substitution in NS1 minimally affected the structure and lipid composition of hexameric NS1, and further suggested that the *in vivo* attenuation was not due to compromised structural integrity of T209L NS1 protein.

### The attenuated phenotype is partially contributed by circulating soluble T209L NS1

To further investigate the mechanisms involved in the attenuated phenotype observed in mice infected with T209L NS1 DENV, we assessed the role of soluble T209L NS1 protein (T209L sNS1) that circulates in the blood during infection. To do so, a depletion experiment was carried out where mice were treated with NS1 hyper-immune serum. As reported by us before (P. X. Lee et al., 2020), sNS1 depletion did not influence disease progression and severity in mice infected with WT virus (Fig. 3), confirming minimal role of circulating sNS1 in viral pathogenesis in this mouse model. In contrast, sNS1 depletion in T209L NS1 DENV-infected mice significantly worsened disease severity, as evidenced by more pronounced body weight loss (Fig. 3D) and increased clinical scores (Fig. 3E) compared to T209L NS1 DENV-infected mice, which correlated with significantly higher viremia titers at day 4 and 6 p.i. (Fig. 3C). This observation thus indicated that while circulating WT sNS1 did not contribute to the virulent phenotype of D2Y98P, T209L sNS1 contributed to the attenuated phenotype of T209L NS1 DENV mutant.

**Figure 3.**
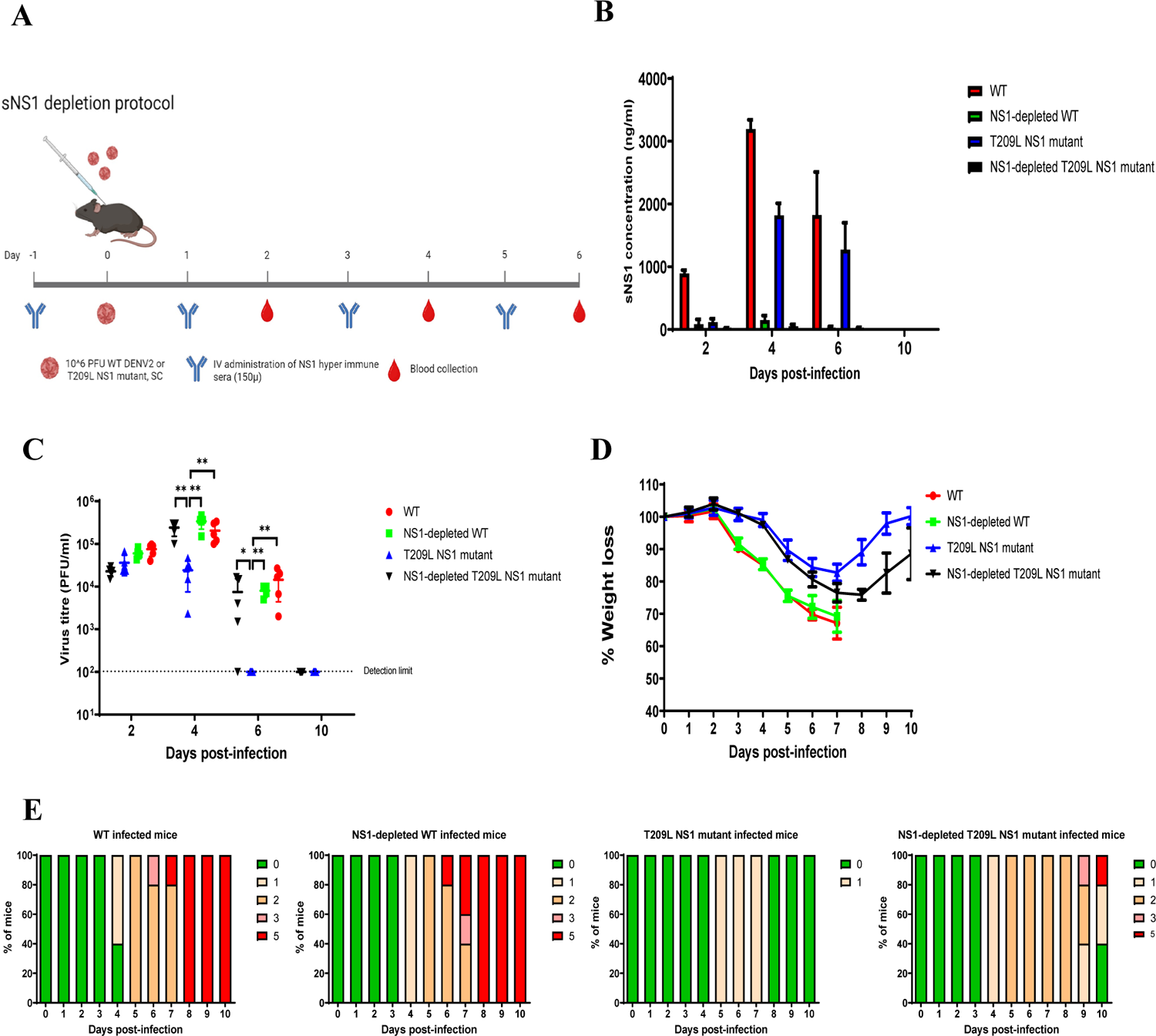
*In vivo* sNS1 depletion. A) Experimental design of sNS1 depletion in infected IFNAR^-/-^ mice. B) Systemic sNS1 concentrations measured by ELISA (n=5). C) Viremia titers measured by plaque assay (n=5). D) Weight loss profile (n=10). E) Clinical scores as described in Legend of Fig. 1 (n=10). Data shown are representative of at least 2 independent biological repeats.

Conversely, exogenous administration of purified T209L sNS1 in WT-infected mice reduced the disease severity and viremia titers at day 6 p.i. compared to WT-infected controls (Fig. 4A-D), further confirming the role of circulating T209L sNS1 in the attenuated phenotype and suggesting that T209L sNS1 had a dominant protective effect over WT sNS1. Of note, exogenous administration of T209L sNS1 did not increase clinical scores and virus titers in T209L-infected mice (Fig. 4A-D), confirming that the attenuated effect was not due to lower circulating sNS1 levels in these mice.

**Figure 4.**
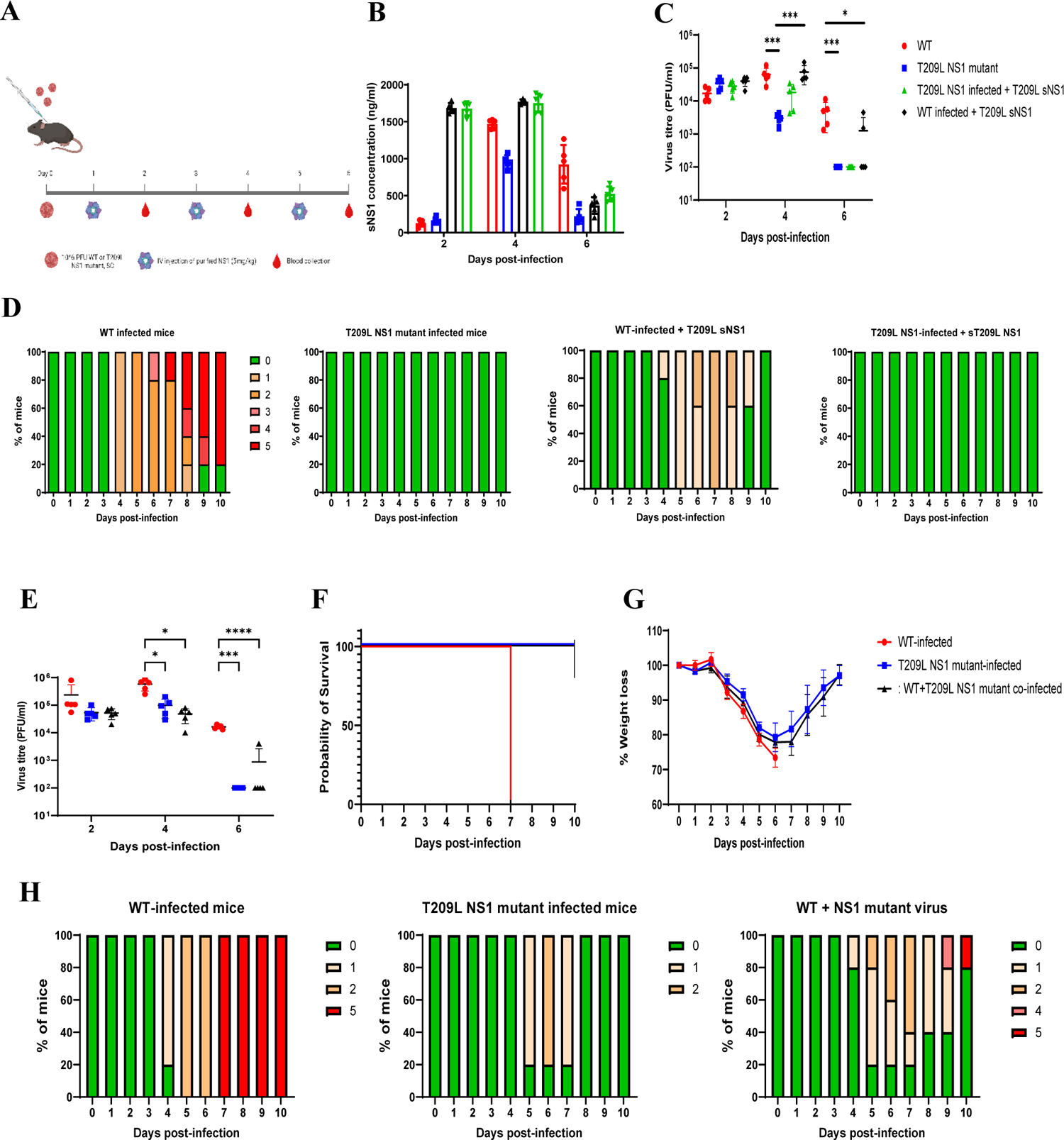
Exogenous administration of sNS1 and co-infection experiments. A-D) Exogenous administration of sNS1 *in vivo.* A) Experimental design. B) Systemic sNS1 concentrations measured by ELISA (n=5). C) Viremia titers measured by plaque assay (n=5). D) Clinical scores as described in legend of Fig. 1 (n=10). E-H) Co-infection with WT DENV and T209L NS1 DENV. IFNAR^-/-^ mice were infected with either WT or T209L mutant (10^6^ PFU per mouse) or were co-infected with WT and T209L NS1 mutant viruses (5 x 10^5^ PFU of each virus). E) Viremia titers were measured by plaque assay (n=5). F) Survival rate (n=10), G) Body weight loss profile (n=10). H) Clinical scores as described in legend of Fig. 1 (n=10). 0 – healthy, 1 – ruffled fur, 2 – hunched back, 3 – lethargy, 4 – limb paralysis, 5 – mice displaying 30% weight loss (euthanasia) Data shown are representative of at least 2 independent biological repeats.

Furthermore, a co-infection experiment was performed where equal amounts of WT and T209L NS1 viruses were co-administered (Fig. 4E-H). Significant disease attenuation was observed in the co-infected mice compared to mice infected with WT DENV only, as evidenced by milder clinical scores, increased survival rate, and lower viremia titers at day 4 and 6 p.i. (Fig. 4). Strikingly, the degree of attenuation observed in the co-infected mice was on par to that observed with mice infected with T209L NS1 DENV only (Fig. 4E-H). Altogether, these data supported that T209L NS1 mutation was dominant over WT NS1 and drives DENV attenuation. Furthermore, while circulating T209L sNS1 contributed to the attenuated phenotype, intracellular T209L NS1 may also play a role.

### Reduced lymphopenia and improved DENV-specific CD8^+^ T cell response in mice infected with T209L NS1 DENV mutant

The data described in the above sections led us to propose a model where T209L NS1 protein (both in circulation and produced intracellularly) modulates some immune function(s) that leads to improved disease outcome. The accelerated viral clearance observed at day 5 p.i. in mice infected with T209L NS1 DENV mutant prompted us to investigate the T cell responses during infection. Indeed, T cells have been reported to play an important role in controlling viral loads, particularly early post-infection when a robust IgG antibody response has yet to be mounted (Lam et al., 2017). We thus analyzed by FACS the T cell subsets present in the blood and spleen at day 5 p.i. Significantly lower total CD4^+^ and CD8^+^ T cell counts were measured in the blood and spleen from infected mice compared to uninfected controls, indicating lymphopenia in both infected groups (Fig. 5A), as previously reported (Potts & Rothman, 2008; Wilder-Smith, Earnest, & Paton, 2004). A consistent trend in lower CD4^+^ and CD8^+^ T cell counts in the WT-infected mice indicated a more pronounced lymphopenia in this group compared to T209L NS1 DENV-infected mice (Fig. 5A). When looking at T-cell subsets, significantly higher levels of naïve CD4^+^ and CD8^+^ T cells, and lower levels of activated CD4^+^ and CD8^+^ T cells were consistently found in mice infected with T209L NS1 DENV mutant compared to WT-infected animals (Fig. 5A).

**Figure 5.**
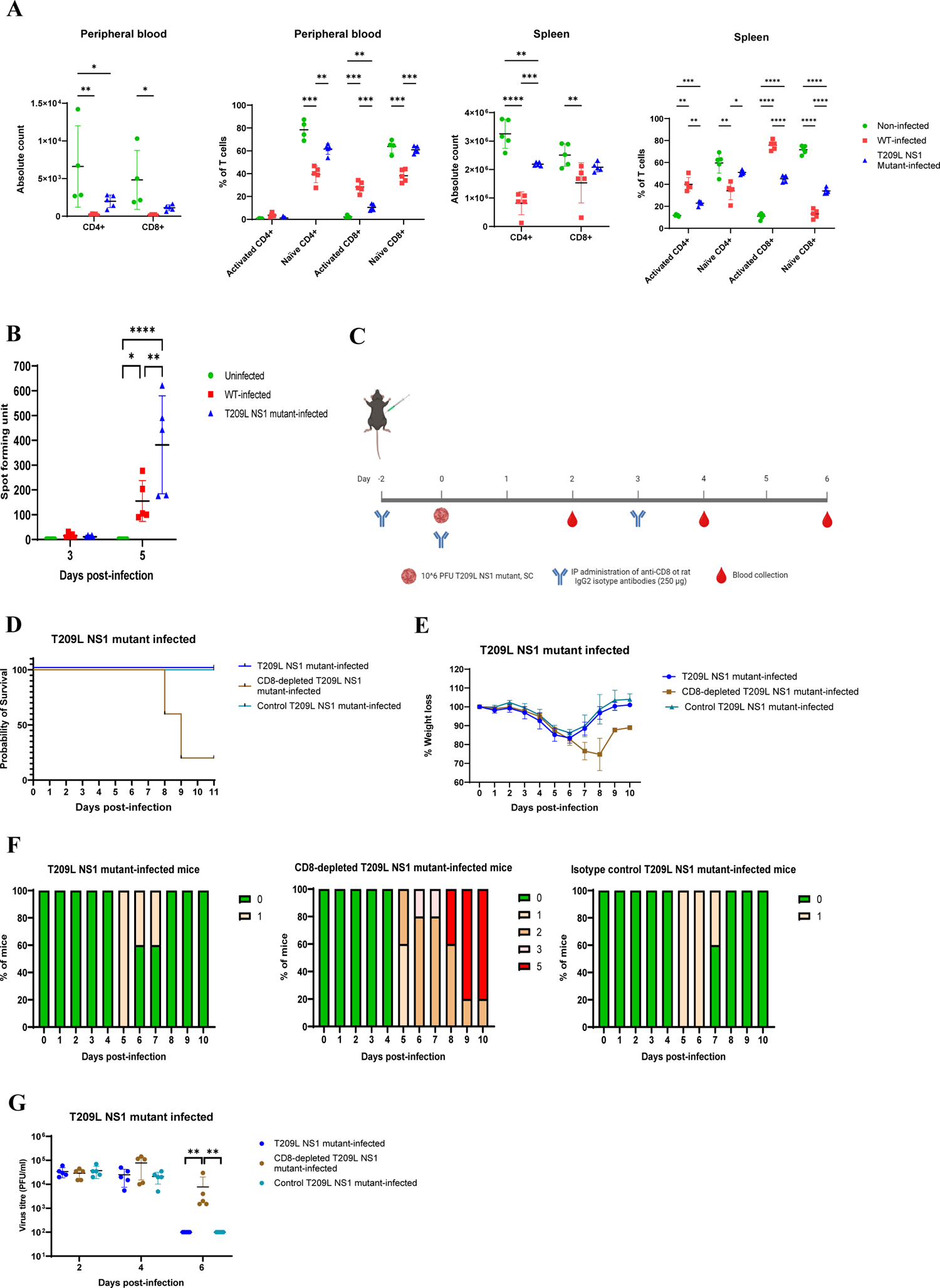
Role of T cells during infection with DENV infection. A) FACS analysis of naïve and activated CD4^+^ and CD8^+^ T cell subsets in WT- or T209L NS1 DENV-infected IFNAR^-/-^ mice at day 5 p.i. B) Splenic CD8^+^ T cells were purified from infected mice at day 3 and 5 p.i. and restimulated with a DENV-specific CD8 immunodominant peptide. IFNγ-ELISPOT was conducted to assess T cell activation. C-G) CD8^+^ T cell depletion experiment in IFNAR^-/-^ mice infected with T209L NS1 mutant. C) Experimental design. D) Survival rate (n=10). E) Body weight loss profile (n=10). F) Clinical scores as described in legend of Fig. 1 (n=10). G) Viremia titers were determined by plaque assay (n=5). Data shown are representative of at least 2 independent biological repeats.

To assess the functionality of the live T cells, CD8^+^ T cells were purified from the spleen of WT- and T209L NS1 DENV-infected mice and were re-stimulated with a DENV-specific immunodominant CD8^+^ T cell epitope (Yauch et al., 2009). Results indicated that at day 5 p.i. the number of IFN-γ producing CD8^+^ T cells was significantly greater in T209L NS1 DENV-infected splenocytes compared to WT-infected splenocytes (Fig. 5B).

Finally, a CD8^+^ T cell depletion experiment was conducted to appreciate the role of these cells during infection with T209L NS1 DENV mutant. CD8^+^ T cell-depleted animals displayed increased disease severity as evidenced by 80% mortality rate and increased clinical scores, which correlated with increased viral titers (Fig. 5C-G), thus supporting a protective role of CD8^+^ T cells in the attenuated phenotype observed in mice infected with T209L NS1 mutant.

Together, these observations suggested that the attenuated phenotype observed in mice infected with T209L NS1 DENV mutant may be linked to greater number of functional CD8^+^ T cells and less severe lymphopenia that contributed to more effective viral clearance.

### T209L NS1 DENV mutant down-modulates the host early inflammatory response

To examine further the mechanisms involved in attenuated disease outcome during infection with T209L NS1 DENV mutant, a comparative bulk RNAseq approach was carried out. Whole blood samples from mice infected with WT or T209L NS1 DENV were harvested at day 3 p.i., a time point at which comparable viral titers were observed in both infected groups (Fig. 1). Cluster analysis revealed that infection with WT NS1 DENV displayed greater innate immune activation and stronger inflammatory response (Fig. 6A) with a pronounced neutrophil activation signature (Fig. 6B), compared to mice infected with T209L NS1 DENV. Consistently, higher systemic levels of TNF-α, IFN-γ, IL-6 and MCP-1 were measured by ELISA at day 3 p.i. in mice infected with WT virus compared to mice infected with T209L NS1 DENV (Fig. 6C). At day 5 p.i. however, comparable cytokine levels between both infected groups were measured, except for IFNγ that was significantly higher in mice infected with T209L mutant, and consistent with increased IFNγ-producing CD8^+^ T cells upon re-stimulation (Fig 4B). This pro-inflammatory cytokine kinetic profile was consistent with vascular leakage in liver and kidneys, which was greater in WT-infected animals at day 4 p.i. but not at day 6 p.i., compared to T209L NS1 DENV-infected group (Suppl. Fig. S3).

**Figure 6.**
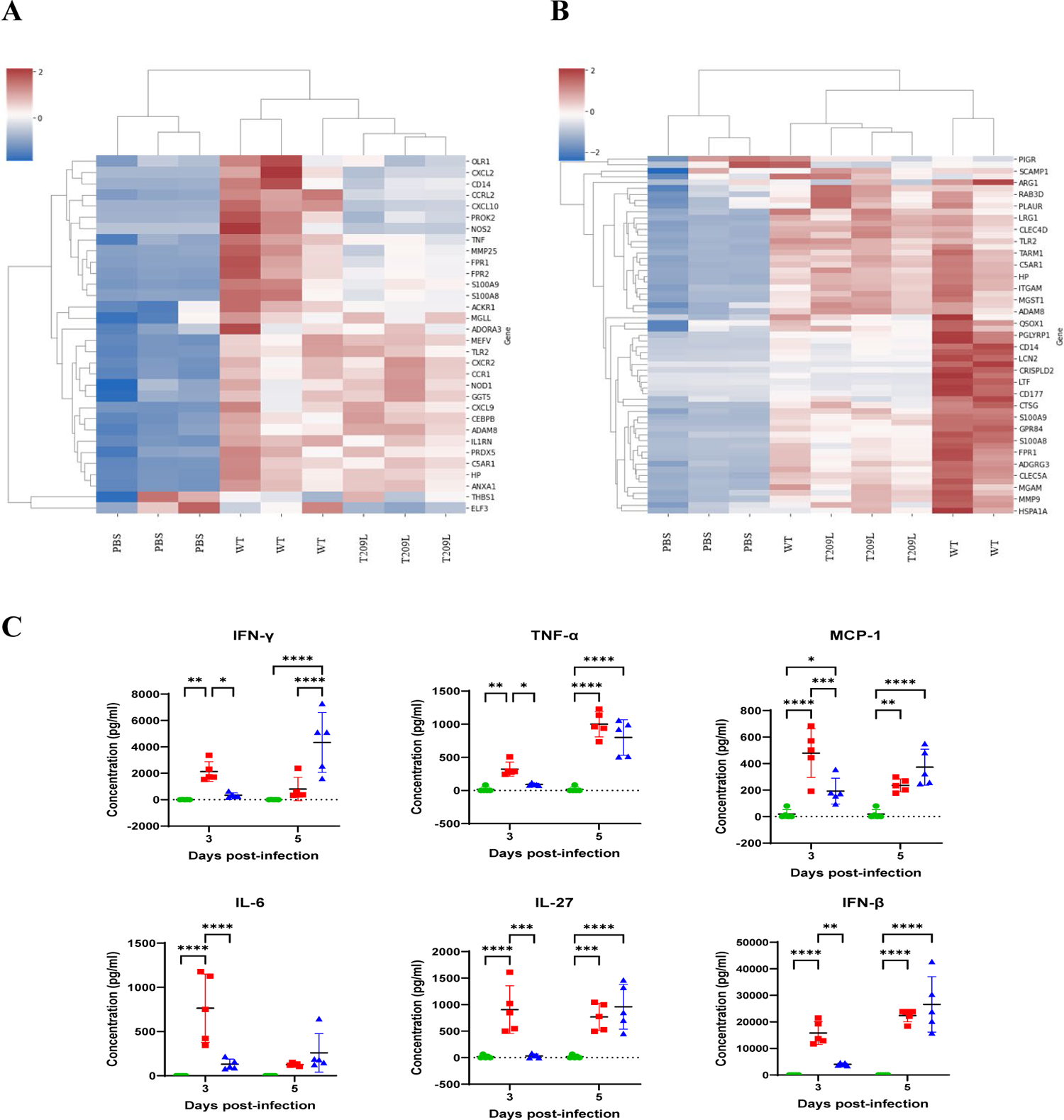
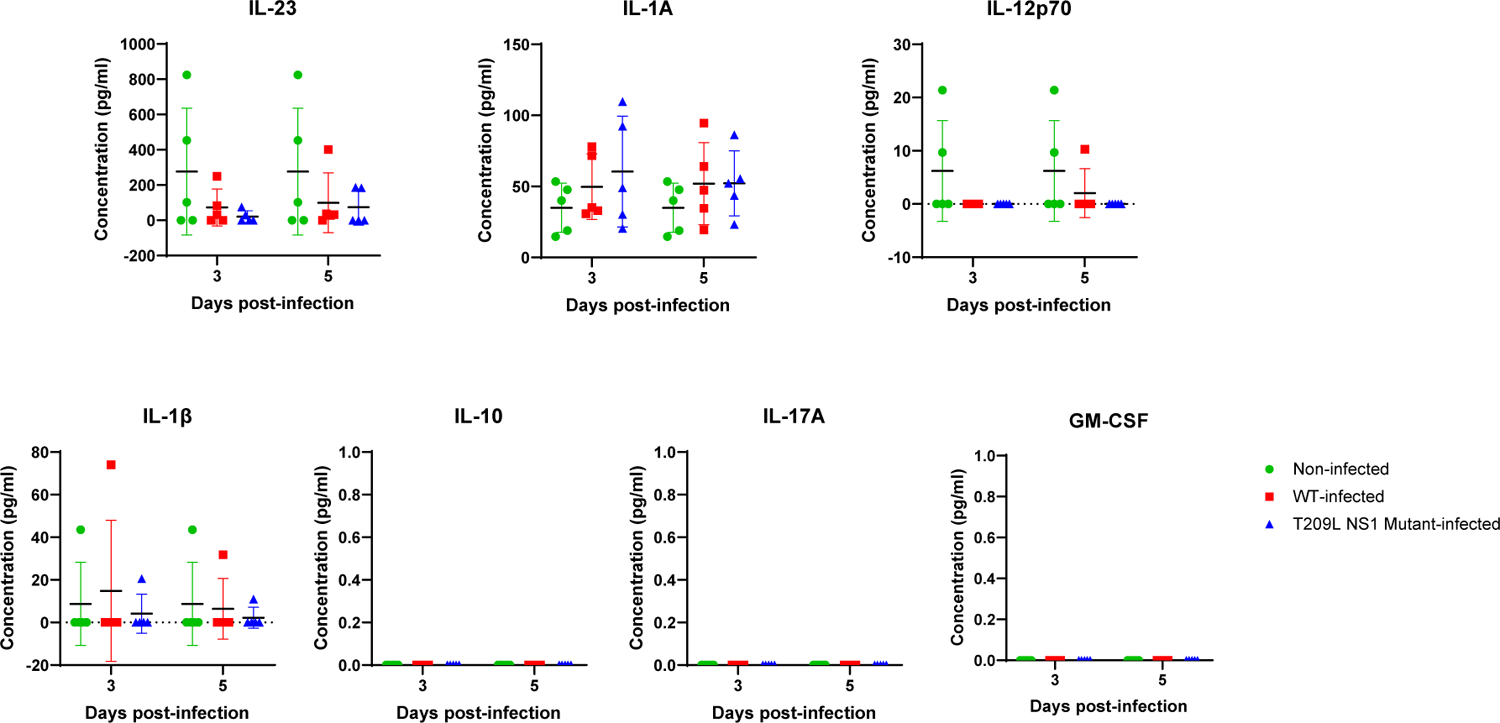
Inflammatory responses in mice infected with WT or T209L NS1 DENV. A-B) Bulk RNA sequencing on whole blood harvested at day 3 p.i. from IFNAR^-/-^ mice infected with WT or T209L NS1 DENV (n=3). A) Innate immune responses and inflammatory responses (Reactome) were attenuated in T209L compared to WT. B) Expression levels of genes related to neutrophil activation and degranulation. C) Systemic cytokine profile measured by ELISA in IFNAR^-/-^ mice infected with WT or T209L NS1 DENV mutant. Data shown are representative of at least 2 independent biological repeats.

In conclusion, both RNAseq analysis and cytokine profile supported the hypothesis that T209L NS1 protein down-modulates the initial host inflammation response during infection, which may in turn influence CD8^+^ T cell activation.

### T209L sNS1 influences CD8^+^ T cell fate via the PD-L1/PD-1 axis

The combined observations of lower early inflammatory cytokine levels, strong down-regulation of genes involved in neutrophil activation, and improved DENV-specific CD8^+^ T cell response in mice infected with T209L NS1 DENV mutant prompted us to probe the involvement of the PD-L1/PD-1 axis in disease severity. Indeed, during infection neutrophils have been shown to engage with T cells through PD-L1/PD-1 interactions, leading to lymphopenia (de Kleijn et al., 2013). Furthermore, studies have reported that PD-L1 expression is modulated by TNF-α, IL-6 and IFN-γ (Bardhan, Anagnostou, & Boussiotis, 2016; de Kleijn et al., 2013; Francisco, Sage, & Sharpe, 2010; Jiang et al., 2019; Wilke et al., 2011). Consistent with this idea, increased absolute counts of PD-L1^+^ activated neutrophils and increased PD-L1 MFI signals were measured in WT-infected mice compared to the T209L NS1 DENV-infected group (Fig. 7A). Furthermore, a trend in increased PD-1 expression was observed on both circulating and splenic activated CD4^+^ and CD8^+^ T cells in WT-infected mice compared to the mutant-infected group, however the differences did not reach statistical significance (Suppl. Fig. S4).

**Figure 7.**
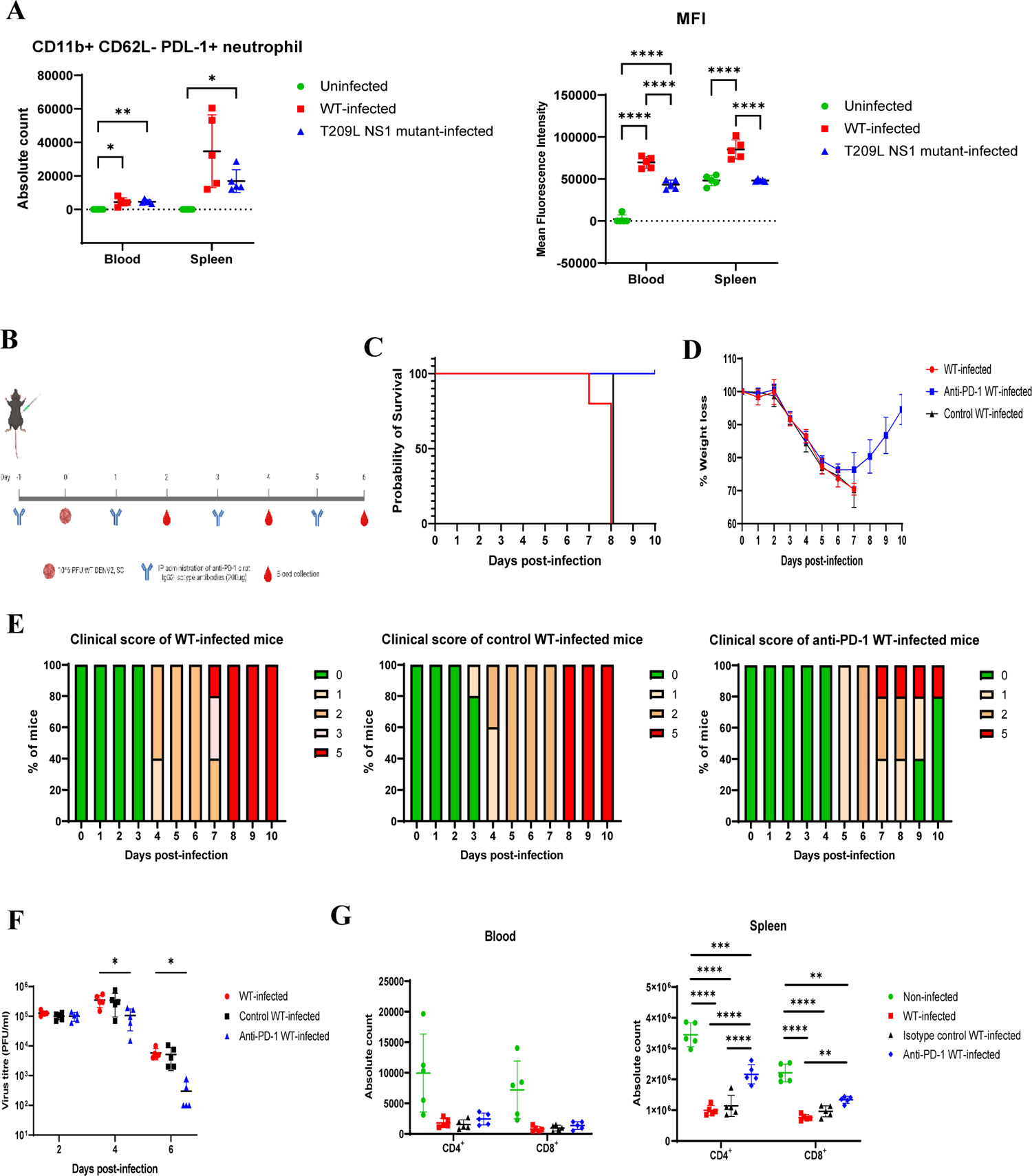
PD-L1 expression on neutrophils and PD-1 blocking experiment. A) FACS analysis of circulating and splenic neutrophils harvested at day 3 p.i. from WT- or T209L NS1 DENV-infected IFNAR^-/-^ mice (n=5). Absolute counts of PD-L1^+^ neutrophils and mean fluorescence intensity (MFI) of PD-L1 expression on activated neutrophils. B-G) PD-1 blockade experiment. B) Experimental design. C) Survival rate (n=10). D) Body weight loss profile (n=8). E) Clinical scores as described in legend of Fig. 1 (n = 10). F) Viremia titers were measured by plaque assay (n=5). G) CD4^+^ and CD8^+^ T cell counts in blood and spleen harvested at day 5 p.i. were analysed by FACS. Data shown are representative of at least 2 independent biological repeats.

To further examine involvement of the PD-L1/PD-1 axis in dengue disease severity, a PD-1 blocking experiment was next performed (Fig. 7B). The results indicated that treatment with PD-1 blocking antibody effectively protected mice from severe disease, as evidenced by 100% survival rate and milder clinical manifestations compared to infected only and isotype treated control groups (Fig. 7C-D). This improved disease outcome correlated with lower viremia titers at day 6 p.i. and increased levels of both CD4^+^ and CD8^+^ T cells in the spleen from PD-1 mAb treated mice (Fig. 7E&F).

These observations thus supported that in WT-infected mice, increased PD-L1 expression on neutrophils may drive T cell death through PD-L1/PD-1 interactions. Consistent with this model, neutrophil depletion in WT-infected mice resulted in attenuated disease severity, as evidenced by improved survival rate, improved clinical scores and lower viremia titers (Suppl. Fig. S5). The partial disease attenuation observed in neutrophil-depleted animals compared to PD-1 blockade however suggested that other immune cells such as Natural Killer cells may also be able to engage T cells via the PD-L1/PD-1 axis.

## DISCUSSION

Previous *in vitro* studies have reported variable effects with flavivirus NS1 de-glycosylated on either N130 or N207 sites. Mutation at N130 on DENV4 NS1 led to reduced viral growth in mammalian and mosquito cells (Pletnev, Bray, & Lai, 1993), whereas the same mutation did not affect viral production of DENV2 16681 strain in mammalian cells (Crabtree et al., 2005). Abrogation of N207 glycosylation site on DENV1 and DENV2 NS1 proteins did not attenuate these viruses in mammalian cells although a delayed cytopathic effect was noted (Crabtree et al., 2005; Tajima et al., 2008); while *in vitro* attenuation was reported with N207 de-glycosylated NGC DENV2 (Pryor & Wright, 1994). Here, we did not observe any significant *in vitro* attenuation when either N-glycosylation site was abrogated on NS1 from D2Y98P DENV2. These inconsistent observations thus suggested that the role of NS1 glycans at N130 and N207 in DENV *in vitro* fitness is highly strain-dependent.

*In vivo*, while N130 mutation has been reported to attenuate DENV2 16681 and DENV4 virulence in suckling mice (Crabtree et al., 2005; Fang et al., 2023; Pletnev et al., 1993), the loss of glycans at this site did not influence D2Y98P virus fitness and virulence in IFNAR^-/-^ adult mice. In contrast, glycans at position N207 were found to play a critical role in the virus virulence, and this observation was consistent with previous studies using another DENV2 strain (Crabtree et al., 2005; Pryor & Wright, 1994). In those earlier studies, the reduced virulence observed was proposed to be linked to the reduced levels of soluble NS1 in circulation. However, our data did not support that the attenuated *in vivo* phenotype observed with T209L DENV mutant was attributable to lower sNS1 levels in circulation. Indeed, a similar reduction in circulating sNS1 levels was measured in mice infected with N130Q mutant, which did not result in attenuation. Instead, several lines of experimental evidence supported that soluble T209L NS1 in circulation contributed to the *in vivo* attenuated phenotype observed. First, antibody-mediated depletion of T209L sNS1 in circulation worsened disease severity in mice infected with T209L DENV mutant, while depletion of WT sNS1 did not attenuate WT infection. Second, exogenous administration of purified T209L sNS1 in mice infected with WT DENV led to significant disease attenuation. Furthermore, co-infection with WT and T209L DENV mutant phenocopied the attenuated phenotype seen with mice infected with T209L DENV alone. These observations therefore supported that T209L NS1 mutant was dominant over WT NS1, and that the attenuated phenotype is likely contributed by both soluble and cell associated T209L NS1 protein.

We further explored the mechanisms by which T209L NS1 attenuates disease severity. Our data supported that NS1 T209L interfered with the interactions between innate immune cells (neutrophils and others) and T cells through the PD-L1/PD-1 axis, limiting activated T cell apoptosis, hence improving T cell-mediated viral clearance. In support of this model, we first found that lymphopenia was significantly less pronounced in IFNAR^-/-^ mice infected with T209L DENV mutant compared to mice infected with WT DENV. Furthermore, CD8^+^ T cell depletion in mice infected with T209L DENV worsened disease severity, confirming the protective role of CD8^+^ T cells in this infection model and as previously reported by us and others (Lam et al., 2017; Weiskopf et al., 2013; Yauch et al., 2010; Yauch et al., 2009; Zellweger, Eddy, Tang, Miller, & Shresta, 2014; Zompi, Santich, Beatty, & Harris, 2012). Second, a comparative transcriptomic approach revealed that mice infected with WT virus displayed greater innate immune cell activation and stronger inflammatory response with pronounced neutrophil activation and degranulation signature compared to mice infected with T209L DENV. Neutrophil activation and degranulation has previously been reported in dengue (Juffrie et al., 2000). Separately, activated neutrophils have been known to engage with T cells through the PD-L1/PD-1 axis during infection, which triggers T cell apoptosis and leads to lymphopenia (de Kleijn et al., 2013). It has also been shown that PD-1-mediated T-cell ligation to PD-L1 alters immunity against viruses by preventing T cell generation and expansion (Bardhan et al., 2016), and subsequently promoting T cell death (Ostrand-Rosenberg, Horn, & Haile, 2014), although this phenomenon had not been described for DENV so far. PD-L1 expression is modulated by IL-6, IFN-γ, and TNF-α (Bardhan et al., 2016; de Kleijn et al., 2013; Francisco et al., 2010; Jiang et al., 2019; Wilke et al., 2011). Consistently, we measured higher PD-L1^+^ neutrophils counts and higher PD-L1 expression on neutrophils in WT-infected mice compared to mice infected with T209L DENV, which correlated with higher systemic levels of IFN-γ, IL-6 and TNF-α early during infection (day 3 p.i.) in mice infected with WT DENV. Involvement of the PD-L1/PD-1 axis in dengue disease severity was directly demonstrated by treating WT-infected mice with an anti-PD-1 blocking mAb, which resulted in complete protection from lethal outcome, accelerated viral clearance and reduced lymphopenia. However, neutrophil depletion only partially attenuated dengue disease in WT-infected mice, and this is likely due to the fact that other immune cells including NK cells, myeloid cells and epithelial cells also up-regulate PD-L1 in response to IFN-γ and TNF-α, and can therefore promote T cell apoptosis through the PD-L1/PD-1 axis (Francisco et al., 2010; Garcia-Diaz et al., 2017; Jiang et al., 2019; Sun, Mezzadra, & Schumacher, 2018). Our data hence support a model whereby de-glycosylated NS1 at N207 interferes with PD-L1/PD-1 interactions, thereby reducing T cell apoptosis and improving viral clearance. One could speculate that such interference involves the down-modulation of key cytokines such as IL-6, IFN-γ and TNF-α early during infection, that limits PD-L1 up-regulation on a variety of immune and non-immune cells. However, the molecular mechanisms by which T209L NS1 exerts this down-modulatory effect remain to be investigated. Importantly, this model implies that in the context of WT DENV infection, glycans at N207 prevent NS1 from exerting this down-modulatory function.

In conclusion, our study has unraveled a novel immune evasion strategy by DENV through NS1 glycosylation. The highly conserved N-glycosylation sites on NS1 across flaviviruses strongly suggest an important role in viral fitness, and mutating these sites should be considered for the development of live attenuated DENV vaccine strains. Furthermore and importantly, our work supported that targeting the PD-1/PD-L1 axis could represent a promising host targeted therapeutic approach to limit T cell death and improve viral clearance in dengue patients.

## MATERIALS AND METHODS

### Ethics statement

All animal experiments were carried out in accordance with the guidelines of the National Advisory Committee for Laboratory Animal Research. Animal facilities are licensed by the regulatory body Agri-Food and Veterinary Authority of Singapore. The described animal experiments were approved by the Institutional Animal Care and Use Committee from National University of Singapore (NUS) under protocol numbers R16-0422 and R18-1400.

### Cell lines and viruses

C6/36 *Aedes albopictus* cell line (ATCC; CRL-1660) was maintained in Leibotvitz’s L-15 medium (Gibco) supplemented with 10% FBS (Gibco) at 28 °C. Baby hamster kidney-21 (BHK-21; ATCC; CCL-10) cell line was maintained in RPMI 1640 (Gibco) supplemented with 10% FBS and cultured at 37 °C with 5% CO_2_. African green monkey kidney epithelial (Vero; ATCC; CCL-81) cell line was maintained in DMEM (Gibco) supplemented with 10% FBS and cultured at 37 °C with 5 % CO_2_. DENV2 (Dengue D2Y98P-PP1; GenBank accession no. JF327392.1) derives from a 1998 Singapore clinical isolate that had been exclusively passaged in C6/36 cells and plaque purified twice in BHK-21 cells, before an infectious clone was made. DENV stock was propagated in C6/36 cell line maintained in Leibovitz’s L-15 medium supplemented with 2 % FBS as previously described (P. X. Lee et al., 2020). Harvested culture supernatants containing the virus particles were stored at −80 °C. Virus titers were determined by plaque assay in BHK-21 cells as described below.

### DENV infection of cell lines

C6/36, BHK-21, and Vero cells were infected at a multiplicity of infection (MOI) of 0.1 with WT D2Y98P, N130Q NS1 mutant or T209L NS1 mutant. Plates were incubated at 37 °C for 1 h with rocking every 15 min for viral adsorption. Each well was rinsed twice with PBS before addition of 200 µl of respective media containing 2 % FBS. The plates were incubated for 4 days at 28^0^C (C6/36) or 37 °C (BHK-21 and Vero), and the culture supernatants were collected at indicated time points post-infection (p.i.). Viral quantification was performed by plaque assay.

### Virus quantification by plaque assay

Virus titer was quantified by plaque assay in BHK-21 cells as previously described (P. X. Lee et al., 2020). Briefly, 45,000 cells/well were seeded in 24-well plates (Nunc) 1 day before plaque assay. Cell monolayers were then infected with 10-fold serially diluted viral suspensions in RPMI 1640 supplemented with 2 % FBS. After 1-h incubation at 37 °C with CO_2_, overlay medium [RPMI 1640 containing 1 % (wt/v) carboxymethyl cellulose and 2 % FBS] was added to each well. After incubation for 4 days at 37 °C with CO_2_, cells were fixed with 4% paraformaldehyde (Sigma-Aldrich) and stained with 0.05 % crystal violet (Sigma-Aldrich). Plaques were counted, adjusted by dilution, and expressed as the number of plaque forming units per milliliter (PFU/mL).

### Generation of de-glycosylated NS1 DENV mutants

D2Y98P RNA genome was extracted using QIAamp Viral RNA Kits (Qiagen). Complementary DNA (cDNA) synthesis was performed using GoScript™ Reverse Transcriptase (Promega) as per manufacturer’s instructions. Four PCR fragments of around 2,700 nucleotides long were generated from cDNA using primer pairs (Suppl. Table S2) with NEB Q5 Hot-Start high-fidelity 2× Master Mix (New England Biolabs). Fragments were gel-purified with MinElute gel extraction Kit (Qiagen) after agarose gel electrophoresis and blunt end cloning was performed into pCR™-Blunt II-TOPO™ Vector (Invitrogen). Site-directed mutagenesis was performed on plasmid that carried E gene to introduce the desired mutation using primer pairs (Table S2). The mutated gene was amplified with other genes fragments via polymerase chain reaction (PCR) and seamlessly assembled with vector containing cytomegalovirus (CMV) promoter sequence, hepatitis delta virus (HDV) ribozyme and simian virus 40 (SV40) poly-A sequence using NEBuilder HiFi DNA Assembly Master Mix (New England Biolabs) at 50 °C for 60 min. This CMV-DENV genome-HDVr-SV40pA assembled product was transfected into BHK-21 cells using Lipofectamine 2000 (Invitrogen). Four to six days later, the viral supernatant was collected and sequenced using Sanger sequencing.

### Mouse infection with DENV

5–6-wk-old IFNAR^-/-^ mice were infected subcutaneously (sc.) with 10^6^ PFU of parental (D2Y98P) or NS1 mutants (N130Q NS1 or T209L NS1) viruses. Mice were monitored daily for body weight loss and for clinical manifestations. Mice were promptly euthanized when they lost 30% of the body weight measured at the start of the infection experiment. In the co-infection experiment, equal amounts (5 x 10^5^ PFU) of WT and T209L NS1 DENV mutant were mixed and administered sc.

### sNS1 depletion in vivo experiment

5–6-wk-old IFNAR^-/-^ mice were immunized intraperitoneally (ip.) three times (weeks 0, 2, and 6) with 20 µg of purified hexameric NS1 from DENV2 D2Y98P strain (custom-made, The Native Antigen Company) or OVA protein (InvivoGen) adjuvanted with 1 µg of monophosphoryl lipid A (MPLA; InvivoGen) and 1:1 volume AddaVax (InvivoGen) as described previously (Beatty et al., 2015). NS1 immune serum was collected 2 weeks after the third immunization and heat inactivated at 56 °C for 30 min before storage at −80 °C. Antibody titers of immune serum were quantified by ELISA. This NS1 hyper immune serum was used for passive transfer experiment into IFNAR^-/-^ mice. One day after ip. administration of NS1 immune serum (150 µl per mouse), IFNAR^-/-^ mice were sc. infected with 10^6^ PFU of DENV2 (D2Y98P or T209L NS1 mutant). A second and third administration of NS1 hyper immune serum were performed at day 3 and 5 post-infection.

### Exogenous administration of NS1 protein to DENV-infected mice

5–6-wk-old IFNAR^-/-^ mice were infected sc. with 10^6^ PFU of D2Y98P or T209L NS1 D2Y98P mutant. 5 mg/kg of purified NS1(D2Y98P or de-glycosylated T209L) (custom-made, The Native Antigen Company) or OVA protein (InvivoGen) was administered intravenously (iv.) to the infected mice at day 1, 3 and 5 p.i.

### Vascular leakage assessment

5–6-wk-old IFNAR^-/-^ mice were infected sc. with 10^6^ PFU of D2Y98P or T209L NS1 D2Y98P mutant. At day 4 or day 6 p.i., Evans blue (10 μl/g; 0.1 % w/v in PBS) was administered intravenously (iv.). Two hours later, mice were euthanized, perfused thoroughly with PBS, and the liver, spleen and kidneys were harvested. To extract the Evans blue from the organs, N,N-dimethylformamide was added to the organs at 4 ml/g of tissue wet weight and incubated overnight at 37 °C. The suspension was then centrifuged at 10,000 rpm for 5 minutes and transferred to 96-well plate. The absorbance was read at OD_620nm_ (Mabtech).

### CD8^+^ T cells depletion in DENV-infected mice

5–6-wk-old IFNAR^-/-^ mice were infected sc. with 10^6^ PFU of T209L NS1 D2Y98P mutant. Anti-CD8 antibody (BioXCell) or rat IgG2b isotype control (BioXCell) were administered ip. (250 μg per dose) to the infected mice 2 days prior to infection, and at day 0 and 3 p.i.

### PD-1 immune blockade

5–6-wk-old IFNAR^-/-^ mice were infected sc. with 10^6^ PFU of D2Y98P or T209L NS1 D2Y98P mutant. 200μg of anti-PD1 antibody (BioXCell) or rat IgG2b isotype control (BioXCell) was administered ip. to the infected mice one day prior to infection, and at day 1, 3 and 5 p.i.

#### ELISAs

Systemic levels of NS1-specific IgG were quantified via indirect ELISA. Purified NS1 protein from D2Y98P (10 ng/well) diluted in PBS was coated onto 96-well enzyme immunoassay plates (Corning Costar) overnight at 4 °C. Plates were washed three times with wash buffer (0.05 % Tween 20 in PBS) and blocked with reagent diluent (2% BSA in wash buffer) for 1 h at 37 °C. Serially diluted serum samples were added to the wells and incubated at 37 °C. Plates were washed three times before the addition of HRP-conjugated anti-mouse IgG (H+L; Bio-Rad; 170-6516) at 1:3,000 and anti-mouse IgG2b (ab97250) at 1:10,000. Plates were incubated for 1 h at 37 °C. After the final three washes, detection was performed by the addition of *o*-phenylene-diamine dihydrochloride substrate SigmaFast (Sigma-Aldrich) and incubated for 30 min at room temperature. The reaction was stopped upon the addition of 2 N H_2_SO_4_. Absorbance was read at 490 nm, and antibody titers were determined by nonlinear regression as the reciprocal of the highest serum dilution with absorbance corresponding to three times the absorbance of blank wells.

Systemic levels of soluble NS1 (sNS1) in mice were quantified via sandwich ELISA as described previously (P. X. Lee et al., 2020). Mouse anti-NS1 Mab62.1 (a kind gift from Profs Subhash Vasudevan, Duke-NUS, and Christiane Ruedl, Nanyang Technological University; 0.1 µg/well) diluted in PBS was coated onto 96-well enzyme immunoassay plates overnight at 4°C. After washing and blocking as described above, diluted serum samples (from 1:100 to 1:2,500) were added to the wells and incubated for 2.5 h at 37 °C. A standard curve was established by twofold serial dilution of recombinant NS1 (D2Y98P) from 25 to 0.39 ng/ml. Plates were washed five times before the addition of HRP-conjugated mouse anti-NS1 Mab56.2 (a kind gift from Profs Subhash Vasudevan and Christiane Ruedl; 25 ng/well), made by conjugating HRP to Mab56.2 using NH_2_ peroxidase labeling kit (Abnova) and incubated for 1.5 h at room temperature. After the final washes, detection was performed with the addition of tetramethylbenzidine (R&D Systems) for 30 min at room temperature. The reaction was stopped with 2 N H_2_SO_4_. Absorbance was read at 450 nm. The concentration of NS1 was calculated based on the standard curve and expressed as the concentration of NS1 in nanograms per milliliter (ng/mL).

#### Detection of soluble inflammatory mediators

Levels of cytokines and soluble mediators present in serum of infected mice were quantified using LEGENDplex Mouse Inflammatory Panel kit (BioLegend) according to the manufacturer’s protocol. Samples were run using Attune Nxt flow cytometer and analyzed using FlowJo software.

#### Flow cytometry

Anti-CD45-BUV395, anti-CD4-BUV805, anti-CD8α-BV786, anti-CD44-BUV737 and anti-CD62L-BV605 were purchased from BD Biosciences. Anti-Ly6G-PE, anti-CD11b-BV650, anti-PD-1-FITC and anti-PD-L1-PE-Cy7 were purchased from Biolegend. Red blood cells (RBC) were lysed using RBC lysis buffer. For surface staining, splenocytes or blood cells were washed and incubated with antibodies for 30 minutes on ice. The cells were then fixed and were read on a Cytek Aurora Spectral Flow Cytometer and analyzed using FlowJo software (Tree Star).

#### CD8^+^ T cells restimulation assay

IFNAR^-/-^ mice were infected sc. with 10^6^ PFU of D2Y98P or T209L NS1 D2Y98P mutant. At Day 5 p.i., spleens were harvested and CD8^+^ T cells were isolated using MACs CD8a (Ly-2) microbeads (Miltenyi Biotech). CD8^+^ T cells (0.5 x 10^5^ or 1 x 10^5^) were co-cultured with splenic dendritic cells (isolated from WT C57BL/6J) at 1:1 ratio in the presence of 10 µg/ml of NS4B_99-107_ peptide for 20 hours in an ELISPOT plate pre-coated with anti-mouse IFNγ. After 20 hours, cells were discarded and biotinylated detection antibody, enzyme conjugate and chromogenic substrate solution were added sequentially. The number of spot-forming units was enumerated using an ELISPOT plate reader (Mabtech).

#### Native PAGE and Western blot analysis

Purified NS1 proteins (1μg) were separated on 8 % Native PAGE and protein bands were transferred to a nitrocellulose membrane. The nitrocellulose membranes were blocked in TBS buffer 0.1 % Tween 20 containing 5 % nonfat dry milk for 1 hour. NS1 proteins were detected using DENV2 NS1 polyclonal antibody (1:1,000 ThermoFisher Scientific) with overnight incubation at 4 °C, followed by incubation with HRP-conjugated anti-rabbit IgG antibody. Clarity™ Western ECL substrate was used for protein detection and exposed to film.

#### RNASeq analysis

5–6-wk-old IFNAR^-/-^ mice were infected sc. with 10^6^ PFU of D2Y98P or T209L NS1 D2Y98P mutant. Blood was collected at Day 3 p.i. and centrifuged at 6,000 *g* for 10 minutes. Red blood cells were then collected and subjected to lysis. After lysis, cells were washed twice in RPMI (Gibco) supplemented with 10% FBS (Gibco) and centrifuged in between washing. RNA was extracted from the cell pellet by adding 1mL of TRIzol and incubated for 5 minutes at room temperature. Subsequently, 200μL chloroform was added to each sample and mixed vigorously for 15 seconds and incubated for 3 minutes. Mixture was then centrifuged at 12,000 *g* at 4 °C for 15 minutes to collect the aqueous phase of the mixture, which was collected into a new clean tube. 1mL of 100 % isopropanol was then added to the aqueous phase and incubated for 10 minutes at room temperature. Further RNA extraction was then performed using RNeasy mini kit (Qiagen) as per manufacturer’s protocol.

For analysis of RNASeq results, raw counts were analysed using the Partek® Genomics Suite®. Differentially expressed genes (DEGs) were identified by comparing gene expression between T209L NS1 mutant and WT, based on fold-change > 2.0 and false discovery rate adjusted p-value < 0.05 using the Benjamini-Hochberg step-up FDR-controlling procedure. For pathway analysis, the upregulated and downregulated DEGs were analysed against the Gene Ontology Biological Process (GOBP) and Reactome databases using the EnrichR tool (Kuleshov et al., 2016). The DEGs enriched in the respective enriched pathways, namely inflammatory response and neutrophil degranulation pathways were also extracted, normalised using Z-score transformation and the relative expression was plotted on clustergrams using on numpy, pandas and seaborn packages in Python.

#### Glycoproteomics

The samples were treated using the single-pot, solid-phase-enhanced sample preparation (SP3) protocol (Hughes et al., 2019). In brief, 30 µg of NS1 protein were denatured and reduced using 8 M urea/50 mM Ambic and 10 mM DTT at 60 °C for 30 min followed by alkylation with IDA in the dark at room temperature for 30 min. The alkylation reaction was subsequently quenched by the addition of 15 µL of 10 mM DTT and incubation at room temperature for 15 min.

Sera-Mag carboxylated speedbeads (Cytiva, Marlborough, MA) were added to the samples in 1:10 (wt/wt) protein/beads ratio. Acetonitrile was added to the suspension to a final concentration of 70 % ACN (v/v) and the final mixture was then incubated at 24 °C with mixing at 1,200 rpm for 10 min. After 10 min, the magnetic beads were immobilized using a magnetic stand and the supernatant was removed. The magnetic beads were rinsed with 80 % ACN and the beads were immobilized again to discard the washing. This was repeated for another two times before the beads were reconstituted in 50 mM Ambic. Sequencing grade trypsin/lysC (Promega, Madison, WI) was added in 1:20 (wt/wt) enzyme/protein ratio to the beads and incubated at 37 °C for 16 hours. After 16 hours, the beads were centrifuged for 1 min at 20,000 *g*, immobilized on the magnetic stand and the supernatant collected into a clean tube. The supernatant was dried down using a vacuum centrifuge and stored at – 20 °C until the next step of analysis.

For glycopeptide enrichment, 30 μg of the sample after SP3 treatment were reconstituted in 30 µL of LCMS grade water and 10 µL of Sera-Mag carboxylated speedbeads (cat. no. 45152105050250) were added to the sample. Acetonitrile was then added to the mixture to a final concentration of 80 % (v/v). The suspension was incubated at 24 °C with mixing at 1,200 rpm for 15 min. After 15 min, the magnetic beads were immobilized using a magnetic stand and the supernatant was removed. The magnetic beads were rinsed thrice with 80 % ACN. LCMS grade water was added to the beads and incubated at 24 °C with mixing at 1,200 rpm for 15 min to elute the bound glycopeptides. The eluted glycopeptides were collected in a clean tube and dried down. The dried sample was reconstituted in LCMS water and injected into the LCMS for analysis.

18μg of the glycopeptide enriched sample were injected onto a Dionex Ultimate RSLC 3000 nano LC system (Thermo Fisher Scientific, Waltham, MA) coupled to an Orbitrap Fusion Tribrid mass spectrometry system (Thermo Fisher Scientific, Waltham, MA) with an Easy-Spray™ nano source. The samples were trapped on a C18 trap column (Acclaim PepMap100 C18, 5 µm, 300 µm x 5 mm, Thermo Fisher Scientific, Waltham, MA) for 2 min before separation on an Easy-Spray™ C18 nano column (75 µm x 150 mm, 2 µm, Thermo Fisher Scientific, Waltham, MA) with a 107 min gradient at a flowrate of 0.3 µL/min. Solvent A was 0.1 % formic acid in water and solvent B was 0.1 % formic acid in ACN. Mass spectrometry data was acquired in the positive ion data dependent acquisition (DDA) mode. Full scan MS spectra data were acquired in the mass range of m/z 500 to 2000, at an ionization source voltage of 2.2 kV, using the Orbitrap detector with resolution set to 120 K, normalized AGC target at 100 %, maximum injection time of 50 ms. MS/MS data was acquired using a top speed mode of 5 s in the Orbitrap detector at 30 K resolution, with an isolation window of 1.6 m/z, stepped HCD collision energy of 20 %, 30 % and 45 %, normalized AGC target of 200 % and maximum injection time of 100 ms. The criteria for MS/MS to be triggered for the precursor ions were minimum intensity of 5e4, charge states of 2 to 6. Dynamic exclusion was set at a duration of 15 s within a mass tolerance of 15 ppm.

Data analysis was performed using Thermo Proteome Discoverer (v2.2.0.388, Thermo Fisher Scientific, Waltham, MA) integrated with Byonic™ (v3.9.6, Protein Metrics, Cupertino, CA) node. The raw files were first matched against Swiss-Prot proteome database with the targeted NS1 protein sequence included. Software parameters were set as followed: missed cleavages set to two, digestion enzyme set to trypsin (full), precursor mass tolerance at 15 ppm, fragment mass tolerance at 20 ppm, fragmentation type to HCD, with modifications of methionine oxidations and deamidation of asparagine and glutamine set as dynamic and carbamidomethylations of cysteine set as static. N-glycosylation modifications were searched against the N-glycans mammalian database by Byonic™. The false discovery rate (FDR) was set to 1%. A focused database with decoys was created from the initial analysis and the focused database was used subsequently for all samples analysed. For samples that have mutations at a particular amino acid, the amino acid mutation was included as part of the modification parameters and set to dynamic. Quantification of glycopeptides was based on peak area performed using the Minora feature detector and precursor ions quantifier nodes in Thermo Proteome Discoverer.

### Lipidomics analysis

Purified NS1 protein samples (50 µL) were mixed with 450 µL 1-butanol/methanol (BuMe, 1:1 v/v) containing lipid internal standards and vortexed for 30 s, followed by 30 min sonication at 20 °C. The samples were centrifuged at 14,000 *g* for 10 min at 10°C. A volume of 400 µL of the supernatant was carefully transferred into vials and dried down completely using a vacuum concentrator system, then resuspended in 100 µL 1-butanol/methanol (BuMe, 1:1 v/v) before MS analysis.

The analysis was carried out on an Agilent 6495 QQQ and Infinity-II LC-MS system, using a Zorbax RRHD Eclipse Plus C18, 95Å (2.1 x 50 mm, 1.8 µm) column. The mobile phases consisted of (A) 10 mmol/L ammonium formate and formic acid (0.1 %) in water/acetonitrile (60:40, v/v) and (B) 10 mmol/L ammonium formate and formic acid (0.1 %) in 2-propanol/acetonitrile (90:10, v/v). Using a flow rate of 0.4 mL/min, the gradient started from 20 % B to 60 % B in 2 min, 60 to 100 % B in 10 min, where it was maintained for 2 min, then the column re-equilibrated at 20 % B for 1.8 min prior to the next injection. The injection volume was 2 µL. Autosampler and column thermostat temperature were at 15 °C and 45 °C, respectively. Total method run time was 15.8 min.

The mass spectrometer conditions were as follows: Capillary voltage 3000 V, drying and sheet gas temperatures were 250 °C, drying and sheet gas flow rates were 14 L/min and 17 L/min, respectively and the nebulizer nitrogen gas flow was set to 35 psi. A semi-targeted analysis, covering the most abundant lipid classes in mammals, was performed in dynamic MRM positive ion mode, using “unit” resolution (0.7 amu) for Q1 and Q3 isolation width. Chromatographic peaks were annotated based on retention time and specific MRM transitions using Agilent MassHunter Quantitative Analysis software (version B.12.0). Internal standards were used to normalize the raw peak areas in the corresponding lipid class and the final lipid concentrations were expressed as nanomole of lipid per g of protein (NS1 protein in the samples).

### Molecular Dynamics Simulations

It was previously estimated that ∼70 lipid molecules can be extracted from NS1 hexamer (Gutsche et al., 2011). D2Y98P strain experimental lipidomics data were used and only lipids with their count greater than 1 % of total lipids were considered. This corresponded to PC:PE:PI:SM with 11:3:83:3 ratio respectively. In total 8 POPC, 2 POPE, 58 SLPI and 2 PSM lipid were randomly placed in a cubic box of 15×15×15 nm (Jo, Kim, Iyer, & Im, 2008). Approximately 100,000 TIP3P water molecules were added to the box and 150 mM NaCl salt whilst neutralizing the overall system charge. Energy minimization was performed using the steepest descent (SD) minimization algorithm with a 0.01 nm energy step size. Unrestrained production run was run for 500 ns in the NPT ensemble. The preequilibrated micelle was subsequently used for simulations with NS1 hexamer. The experimental structure (PDB: 4O6B) (Akey et al., 2014) of the NS1 hexamer dengue serotype 2 was used. *In silico* mutations to experimentally studied strain D2Y98P were performed using CHARMM-GUI (Jo et al., 2008). Protein charges were assigned according to neutral pH with charged termini. The wild type (WT) NS1 hexamer N-linked glycan composition at site N130 corresponded to Hex:3 HexNAc:7, while N207 site corresponded to Hex:6 HexNAc:3 according to experimental data (Supplemental Fig. S2). The preequilibrated micelle was placed in the center of the hexamer hydrophobic β-roll domains. All lipids overlapping with protein based on the 0.3 nm cutoff distance were moved away from the hydrophobic domain (residues 1-29). The energy minimization in vacuum was performed using SD algorithm. The WT construct was placed in a dodecahedron box with ∼17 nm box edge. Approximately 90,000 TIP3P water molecules were added to the box and 150 mM NaCl salt whilst neutralizing the overall system charge. Energy minimization was performed using the steepest descent minimization algorithm with a 0.01 nm step size. The system was equilibrated in the NPT ensemble for 200 ns with position restraints on protein alpha carbons and a force constant of 1000 kJ mol^-1^ nm^-2^. The equilibrated i) WT-lipid complex was used to generate mutation T209L, which resulted in loss of glycans at position N207. Both systems were subjected to a production run of 500 ns in the NPT ensemble which involved weak position restraints on protein alpha carbons with a force constant of 100 kJ mol^-1^ nm^-2^. All simulations were performed using the GROMACS 2018.3 simulation package (Van Der Spoel et al., 2005) utilizing the CHARMM36m (Huang et al., 2017) force field with the TIP3P water model (Jorgensen, Chandrasekhar, Madura, Impey, & Klein, 1983). A temperature of 310 K was maintained using the velocity rescaling thermostat with an additional stochastic term using a time constant of 1 ps. Pressure was maintained semi-isotropically at 1 atm using the Parrinello-Rahman barostat (Parrinello & Rahman, 1981) and a time constant of 5 ps. All bonds which involved hydrogens were constrained using the LINCS algorithm. Equations of motion were integrated using the leap-frog algorithm with a time step of 2 fs. Long-range electrostatic interactions were described using the particle mesh Ewald method (Essmann et al., 1995). The short-range electrostatics reals pace cut-off was 1.2 nm and the short-range van der Waals cut-off was also 1.2 nm. Periodic boundaries conditions were applied in all directions. Simulations were performed on: i) an in-house Linux cluster composed of 8 nodes containing 2 GPUs (Nvidia GeForce RTX 2080 Ti) and 24 CPUs (Intel® Xeon® Gold 5118 CPU @ 2.3 GHz) each as well as on ii) National Supercomputing Center (https://www.nscc.sg) using 2 nodes containing 128 cores and 256 logical cores (AMD EPYC™ 7713 @ 2.0 GHz) each. All simulations snapshots were generated using VMD (Humphrey, Dalke, & Schulten, 1996). The glycan-lipid contacts were calculated based on the 0.3 nm cutoff distance and were averaged over last 100 ns of trajectory. The buried solvent accessible surface area (SASA) between NS1:glycans and lipid cargo was calculated as a sum of NS1:glycans and lipids before subtracting (TLR4:glycan)–lipid SASA.

### Statistical analyses

Data analyses were performed using Graphpad Prism 9.0. Statistical comparison was conducted using nonparametric Mann–Whitney *U* test or two-way Anova and Sidak’s multiple comparison test. Comparison of survival rates was performed using log-rank (Mantel–Cox) test. Differences were considered significant (*) at P < 0.05.

