## Supplemental Figures for "De-glycosylated non-structural protein 1 enhances dengue virus clearance by limiting PD-L1/PD-1 mediated T cell apoptosis"

**Suppl. Table S1. De-glycosylated NS1 mutants generated by site-directed mutagenesis.** Mutation stability was assessed after 3 consecutive passages in C6/36 cell line.

| Mutants | Amino acid substitution | Change in amino acid chemical property | Features |
| --- | --- | --- | --- |
| N130Q | Asn to Glu | No change | Stable after 3 passages |
| N207H | Asn to His | Polar to basic | Reversion after 3 passages |
| N207Q | Asn to Glu | No change | Reversion after 3 passages |
| T209A | Thr to Ala | Polar to non-polar | Reversion after 3 passages |
| T209L | Thr to Leu | Polar to non-polar | Stable after 3 passages |
| T209V | Thr to Val | Polar to non-polar | Reversion after 3 passages |

**Suppl. Table S2. Primer pairs used for virus genome amplification and site-directed mutagenesis.**

| Primer Name | Primer Name Primer Sequence (5’ to 3’) |
| --- | --- |
| D2Y98P-F1-FP | CTGGTTTAGTGAACCGTCAGAGTAGTTAGTCTACGTGGAC |
| D2Y98P-F1-RP | CTCACAACGCAACCACTATCGGCCTGCACCATAACTCC |
| D2Y98P-F2-FP | TGGGAGTTATGGTGCAGGCCGATAGTGGTTGCGTTGTG |
| D2Y98P-F2-RP | ATTGCTGGAAGGTATCTCTTTGTTTTTCCTGCTCCTGG |
| D2Y98P-F3-FP | ACCCAGGAGCAGGAAAAACAAAGAGATACCTTCCAGCAATAGTCAGAGAAG |
| D2Y98P-F3-RP | TTTGAAGACGCACCAGATTCCAACCATATGTTGACATGG |
| D2Y98P-F4-FP | CCCATGTCAACATATGGTTGGAATCTGGTGCGTCTTCAAAG |
| D2Y98P-F4-RP | TGGAGATGCCATGCCGACCCAGAACCTGTTGATTCAAC |
| Vector (CMV, HDV ribozyme and SV40 PA) FP | CTGTTGAATCAACAGGTTCTGGGTCGGCATGGCATCTC |
| Vector (CMV, HDV ribozyme and SV40 PA) RP | GTCCACGTAGACTAACTACTCTGACGGTTCACTAAACCAGC |
| 130 N-Q FP | CTCTCCACAGAGCTTCATCAACACACCTTTCTCATTGA |
| 130 N-Q RP | CAATGAGAAAGGTGTGTTGATGAAGCTCTGTGGAGAGCAT |
| 209 T-L FP | CTCAATGACCTATGGAAGATTGAGAAAGCCTC |
| 209 T-L RP | GAGGCTTTCTCAATCTTCCATAGGTCATTGAG |

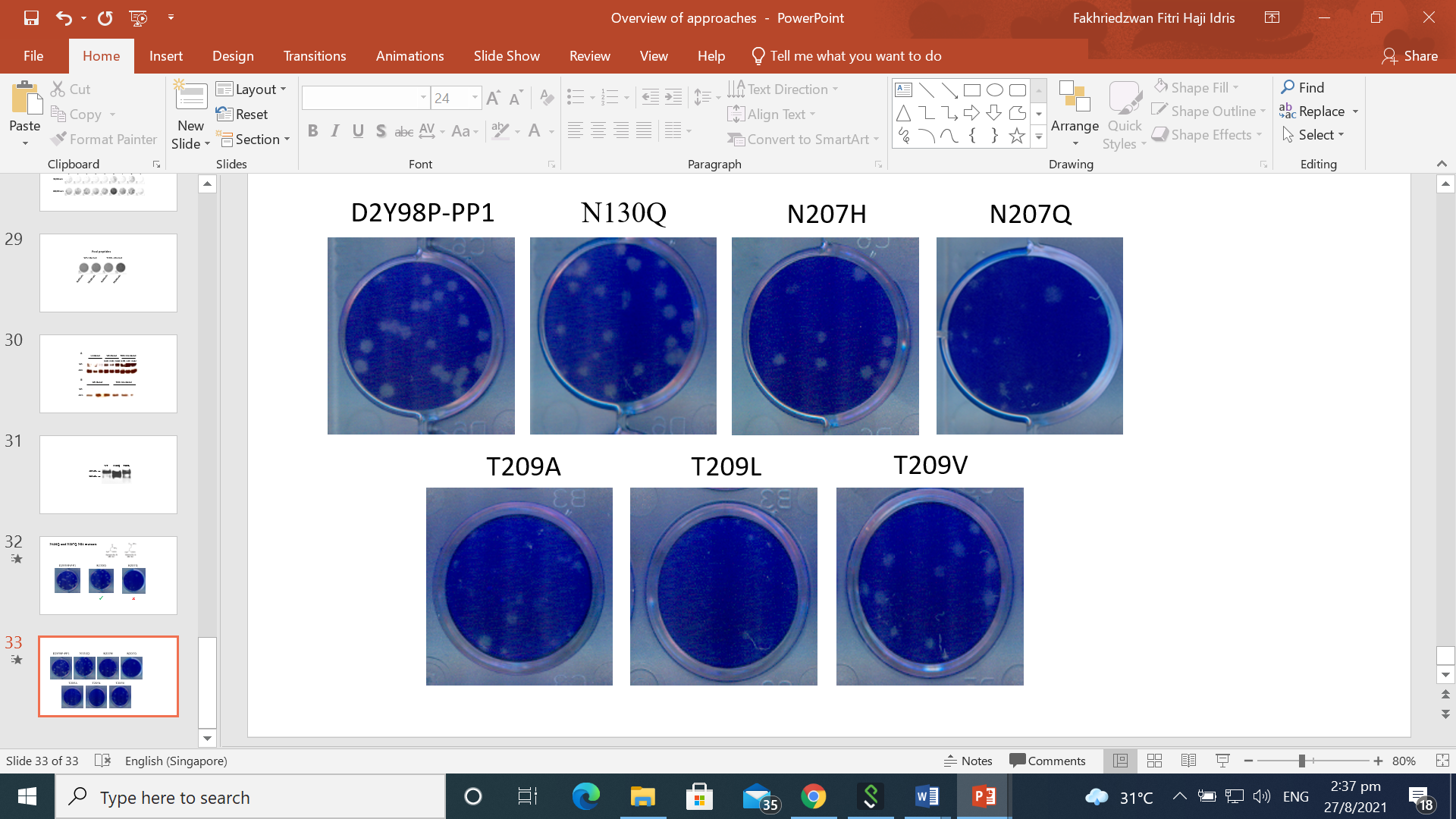

**Supplementary Figure S1. Plaque morphology of WT and de-glycosylated NS1 DENV mutants.**

The viruses were amplified in C6/36 cells and plaqued on BHK-21 cells. Data shown are representative of at least 2 independent biological repeats.

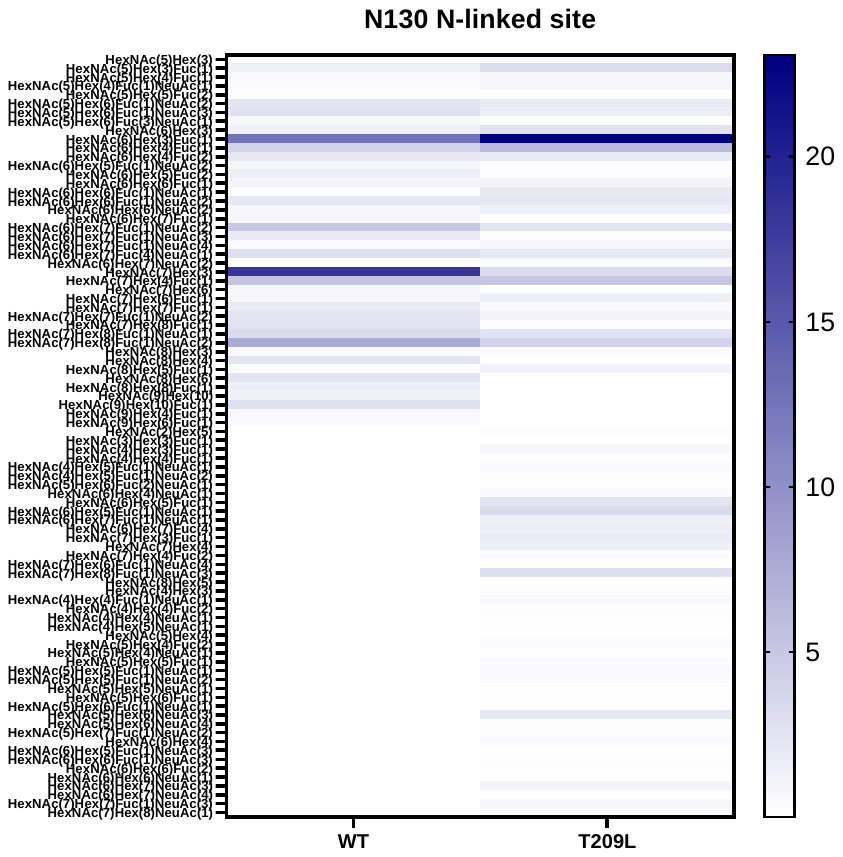

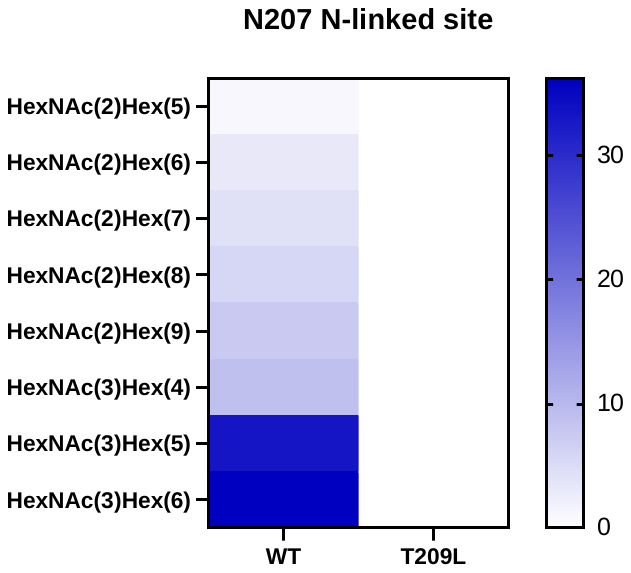

**Supplementary Figure S2. Glycan profiles of WT and T209L sNS1.** Glycomic and glycoproteomic approaches were utilized to determine the glycan species as well as quantifying the abundance of each glycan motif present on both N-glycosylation sites. Data shown are representative of at least 2 independent biological repeats.

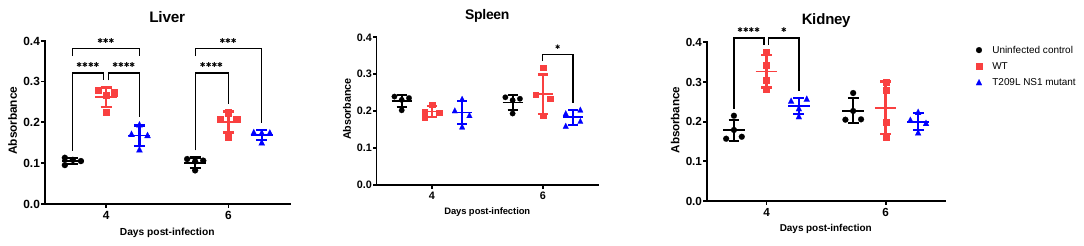

**Supplementary Figure S3. Vascular leakage in mice infected with WT or T209L DENV.** Vascular leakage was determined by Evans Blue assay in mice infected with WT or T209L DENV at day 4 and 6 p.i. Data shown are representative of at least 2 independent biological repeats.

**Blood**

**MFI**

**Proportion of T cells**

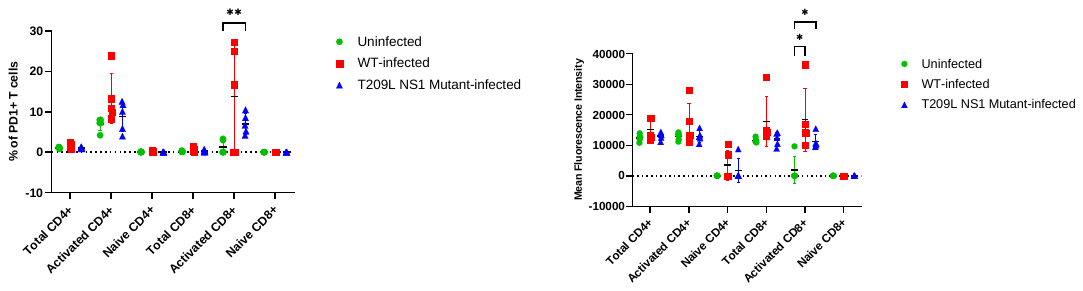

**Spleen**

**MFI**

**Proportion of T cells**

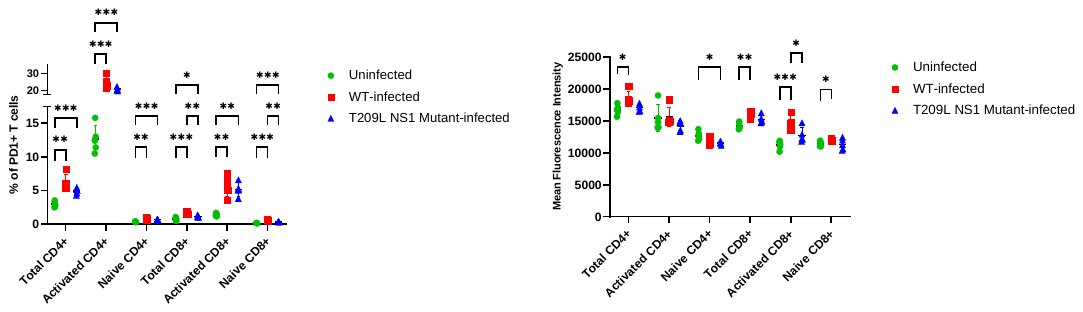

**Supplemental Figure S4:** **PD1 expression on T cells.** FACS analysis of circulating and splenic T cells harvested at day 3 p.i. from WT- or T209L NS1 DENV-infected IFNAR^-/-^ mice (n=5). Percentages of PD-1^+^ T cells and mean fluorescence intensity (MFI) of PD-1 expression on T cells. Data shown are representative of at least 2 independent biological repeats.

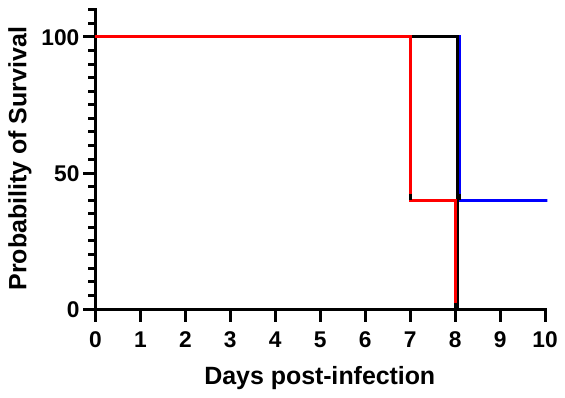

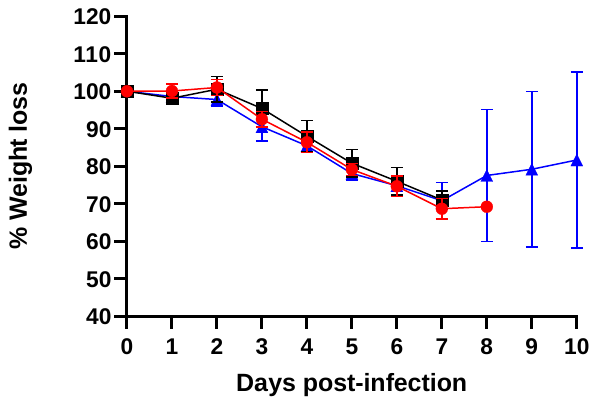

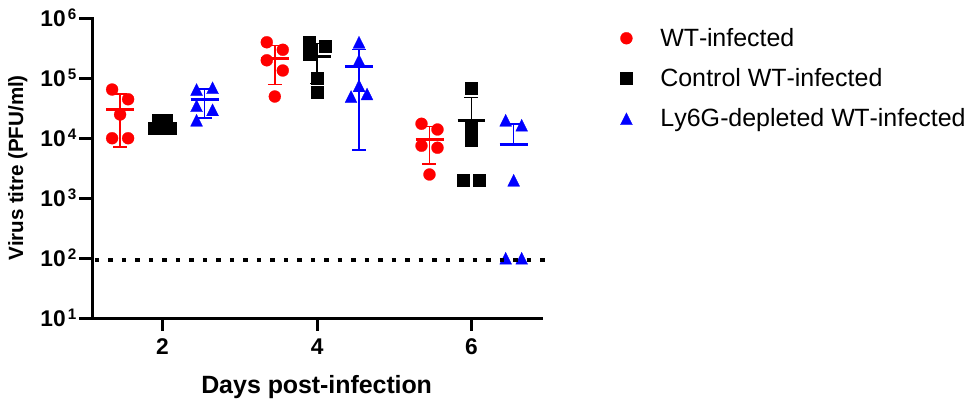

**D**

**C**

**B**

**A**

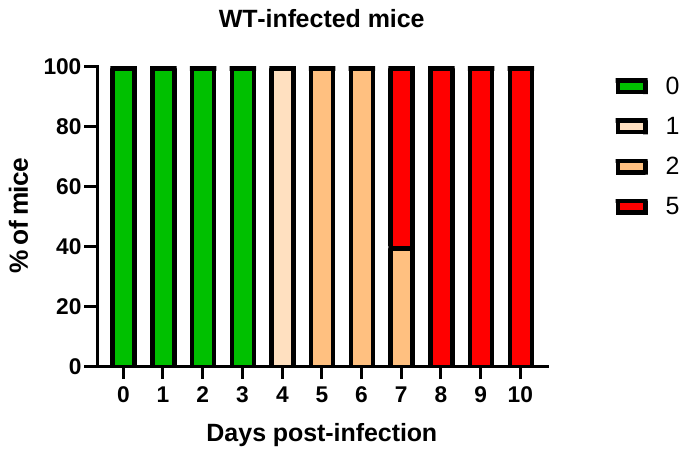

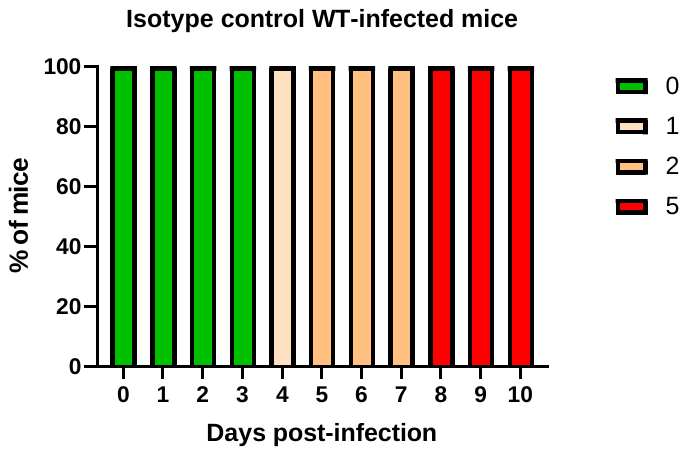

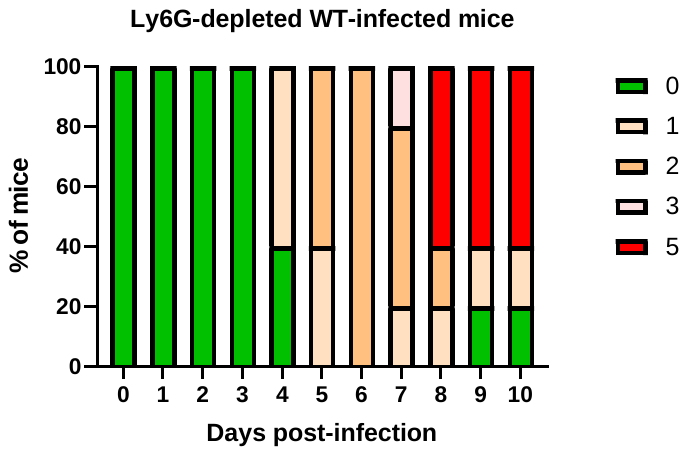

**Supplementary Figure S5. Neutrophil depletion experiment.** A) Survival rate (n=10). B) Body weight loss profile (n=10). C) Viremia titers were determined by plaque assay (n=5). D) Clinical scores (n=10): 0 – healthy, 1 – ruffled fur, 2 – hunched back, 3 – lethargy, 4 – limb paralysis, 5 – mice displaying 30% weight loss (euthanasia). Data shown are representative of at least 2 independent biological repeats.
